# Neuron-like function of the nephron central command

**DOI:** 10.1101/2021.12.06.471478

**Authors:** Georgina Gyarmati, Urvi Nikhil Shroff, Anne Riquier-Brison, Sean D. Stocker, Audrey Izuhara, Sachin Deepak, Yibu Chen, Daniel Biemesderfer, Aaron W. James, Liliana Minichiello, Berislav V. Zlokovic, Janos Peti-Peterdi

## Abstract

Interoceptive neurons that sense and regulate our internal milieu have been identified in several organs except in the kidney cortex despite its major importance in maintaining body homeostasis. Here we report that the chief kidney cell type of the macula densa (MD) forms coordinated neural networks in each nephron that resemble peripheral ganglia. A combined in vivo single-cell 4D physiology (sc4DP) and scRNA sequencing approach identified the MD mechanisms of neuronal differentiation, heterogeneity (pacemaker MD cells), sensing of the local and systemic environment via multi-organ crosstalk, and regulation of organ functions by acting as the nephron central command. Consistent with their neuron-like nature, MD cells express the molecular fingerprint of neurodegeneration. Here we put forth the single-cell MD model and concept of local neural networks that control organ and body functions via interoception in normal physiological state and use an integrated mechanism of neurodegeneration in disease.

## INTRODUCTION

The central and peripheral nervous system senses and integrates information about the inner state of the body through the process of interoception, which is essential for maintaining mental and physical health. Interoception involves a variety of neural and non-neural cells and processes in several internal organs by which an organism senses, interprets, integrates, and regulates signals from within itself (Chen et al., 2021b). There are well-established bidirectional interoceptive communications between the brain and peripheral neural networks in several organs throughout the body including the heart, intestine and bladder (Berntson and Khalsa, 2021; Khalsa et al., 2018). Within the peripheral neural networks, chief hub or pacemaker cells are known to exist and function as the central command for the organs in which they reside, for example the sinoatrial node (heart), islets of Langerhans (pancreas), and the interstitial cells of Cajal (intestine) (Da Silva Xavier and Rutter, 2020; Drumm et al., 2017; Liang et al., 2021).

Despite playing major roles in maintaining body homeostasis, the kidneys have been largely overlooked in the context of interoception. This is largely due to our incomplete knowledge of the neural control mechanisms of the kidney and the kidney-brain axis that is currently limited to the anatomy and function of sympathetic efferent (cell bodies in pre and paravertebral ganglia) and sensory afferent nerves (cell bodies in dorsal root ganglia) and nerve endings in the renal cortex (Osborn et al., 2021). Importantly, no neuronal somata have been described in the renal cortex, although anatomically and functionally far less-complex organs usually have local neural control and neural networks.

One chief, but understudied cell type in the kidney is called macula densa (MD) localized in a strategically central position at the vascular pole entrance of the kidney filter (glomerulus). MD cells are known to provide key physiological control of basic kidney functions including glomerular filtration rate, renal blood flow and renin release (Bell et al., 2003; Peti-Peterdi and Harris, 2010), but the true nature of these cells has been largely unknown due to their inaccessibility. According to the existing paradigm, MD cells can sense alterations in renal tubular fluid variables including NaCl content and metabolic intermediates, that via various intracellular signaling pathways trigger the synthesis and release of chemical mediators. These mediators then in turn act in a paracrine fashion on final effector cells, including renin producing cells to secrete renin, or contractile vascular smooth muscle cells of glomerular arterioles to regulate vascular resistance and organ blood flow (Bell et al., 2003; Peti-Peterdi and Harris, 2010). Importantly, the MD-specific expression of the neuronal type of nitric oxide synthase (*Nos1*) in the renal cortex has been long established (Mundel et al., 1992). The recent microanatomical discovery of long, axon-like MD cell processes (Gyarmati et al., 2021) and preliminary transcriptomic analysis of a limited number of single MD cells expressing neuroepithelial features (Chen et al., 2021a) suggested the neuronal differentiation of MD cells. However, the molecular and functional details of potential neuron-like MD cell functions and their (patho)physiological significance are unknown.

In our quest to better understand the molecular and cellular mechanisms and the role of kidneys in interoception, we established a new comprehensive research toolbox to study MD cells, specifically their neuron-like features in unprecedented detail in the normal physiological and disease states. Here we used this new MD research toolbox to address the hypothesis that MD cells are interoceptive neuron-like cells in the kidney that sense the local and systemic environment, process and send signals to renal and central effectors to maintain homeostasis. Based on the new results, we developed a single-cell type MD model and concept of local neural networks that are applicable to many other organ systems and relevant to modeling kidney and neurodegenerative diseases.

## RESULTS

### Neuron-like functional features of MD cells *in vivo*

Intracellular calcium (Ca^2+^) transients and signaling are classic functional readouts of excitable cells. Therefore, intravital Ca^2+^ imaging using multiphoton microscopy (MPM) was performed in newly established genetic mouse models to directly visualize the activity of MD cells versus all renal cell types in the intact living kidney cortex. We applied three main strategies to study cell physiology in four dimensions (4DP, in tissue volume over time) in either comparative cell (cc4DP), multi-cell (mc4DP), or single-cell (sc4DP) modes as summarized in Fig. S1A. To make comparisons to other renal cell types (in cc4DP mode), Sox2-GT mice were developed with ubiquitous expression of the genetically encoded calcium reporter GCaMP5 (G) along with the calcium-insensitive tdTomato (T) in all kidney cell types. Time-lapse intravital MPM imaging of Sox2-GT mice found robust, spontaneous Ca^2+^ transients in MD cells (Fig. S1B, Supplement Video 1). Among all cells at the glomerular vascular pole including the highly contractile vascular smooth muscle cells of the glomerular arterioles, MD cells showed by far the highest cumulative elevations in Ca^2+^(Fig. 1A). MD cell Ca^2+^ signals did not conduct to adjacent contractile vascular cells as evident from video recordings (Supplement Video 1) and from the lower frequency of MD cell versus contractile mesangial and vascular smooth muscle cell Ca^2+^ transients (Fig. 1A). In addition, the robust MD cell Ca^2+^ transients were autonomous (preserved in freshly isolated single MD cells, Fig. S1C) and unique to MD cells in the renal cortex (based on comparison to other renal tubule epithelial cells (Fig. S1B).

**Figure 1.**
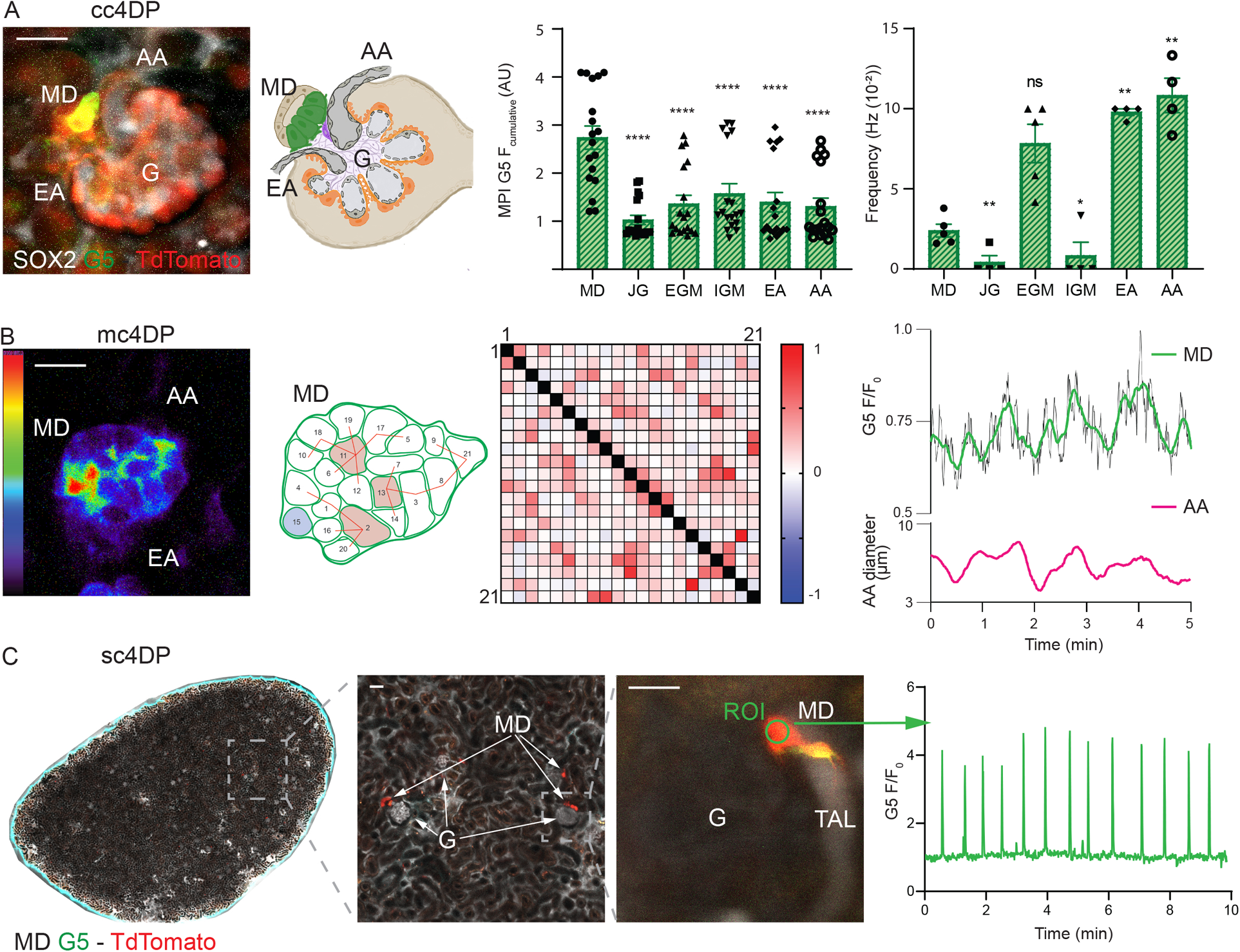
Intravital imaging of macula densa (MD) cell calcium in the intact renal cortex. **(A)** Comparative cell 4D physiology (cc4DP) mode using Sox2-GT mice. Left: Representative maximum projection image (MPI) of a 2-min time-lapse recording (shown in Supplement Video 1) of the Ca^2+^ activity of MD and other renal cell types. Note the highest cumulative Ca^2+^ elevations in MD cells (green cells in the attached anatomical drawing of the same glomerulus). Center: Comparison of GCaMP5 fluorescence intensities (F_cumulative_) in single MD, juxtaglomerular (JG) renin, extra (EGM) and intra-glomerular (IGM) mesangial cells, and afferent (AA) and efferent arteriole (EA) vascular smooth muscle cells. Right: Comparison of the frequencies of Ca^2+^ transients in glomerular vascular pole cell types. *: p<0.05, **: p<0.01, data are mean ± SEM, n=1-4 cells from n=5 mice. **(B)** Multi-cell 4D physiology (mc4DP) mode using Sox2-GT mice. Left: Representative intensity-based pseudocolor MPI image of a horizontal optical section of a whole-MD (shown in Supplement Video 1). Note the heterogeneity of GCaMP5 fluorescence between individual MD cells. Center: Functional cell-to-cell connectivity map of the same MD plaque and horizontal plane showing all 23 individually numbered MD cell positions. Red line connecting individual cell pairs indicates that the strength of the cell pair correlation (Pearson’s R value) was >0.35. Red/blue cell color indicates hub/lone cells, respectively. Heat-map shows Pearson’s coefficient of each cell pair in a two-color gradient (low correlation blue, high correlation red). Right: Representative simultaneous recording of changes in whole-MD GCaMP5 signal and the adjacent AA diameter over time. **(C)** Single-cell 4D physiology (sc4DP) mode using MD-GT mice. Left: Representative tile scan overview image of a 2 × 3 mm surface area of the renal cortex of MD-GT mice. Alexa Fluor 680-conjugated bovine serum albumin was injected iv to label the circulating plasma (greyscale), and the kidney capsule is shown by second harmonic generation (SHG) signal (cyan). A magnified region as shown demonstrates the expression of GT reporters exclusively in MD cells (arrows) at the vascular pole of glomeruli (G). Center: High-power view of a single glomerulus and MD at the end of the thick ascending limb (TAL) tubule segment (tubule fluid in greyscale). Note the long, axon-like basal cell processes of MD cells (also in Supplement Video 1). Right: Representative recording of regular Ca^2+^ firing in a single MD cell. Note the 4-fold elevations in cell Ca^2+^ compared to baseline during Ca^2+^ transients. Bars are 20 μm.

Intravital MPM imaging of the entire MD plaque (formed by ~25 individual MD cells) in horizontal optical sections (in mc4DP mode) revealed considerable cellular heterogeneity, with individual MD cells featuring either low, medium, or high Ca^2+^ activity (Fig. 1B, Supplement Video 1). Several, but not all MD cells in multiple plaque regions showed rapidly and laterally propagating Ca^2+^ responses, further suggesting the heterogeneity of cell-to-cell communication and coordination between individual MD cells (Fig. 1B, Supplement Video 1). However, the robust Ca^2+^ signaling activity remained spatially confined to the MD plaque area and did not propagate to adjacent tubular or vascular segments (Supplemental Video 1). A cell-to-cell connectivity map with preserved MD architecture and a heat map were generated based on Pearson’s correlation analysis of time-lapse Ca^2+^ recordings between all MD cell pairs (Fig. 1B). The functional connectivity map (MD connectome) of this one specific MD plaque identified three “hub” cells within the whole MD that had the highest number of connections, and one “lone” cell that had zero connections (Fig. 1B). Interestingly, the whole-MD Ca^2+^ readout showed clustering of the single-cell Ca^2+^ transients and their regular oscillations over time (Fig. 1B, Fig. S2D). Importantly, the whole-MD Ca^2+^ oscillations were simultaneous with the rhythmic changes in the diameter of the adjacent glomerular afferent arteriole (AA), with the peak MD Ca^2+^ phase-matching the peak AA diameter (vasodilatations) (Fig. 1B). Cumulative single-cell Ca^2+^ elevations and firing frequency showed normal distribution (Fig. S1B).

To study the spatial and dynamic details of MD cell Ca^2+^ responses with truly single-cell resolution (in sc4DP mode), MD-GT mice with MD cell-specific expression of GT were generated with partial tamoxifen induction strategy as used recently for single-cell expression and targeting (Gyarmati et al., 2021). GT reporter expression was entirely specific for MD cells in the MD-GT mouse renal cortex (Fig. 1C), and illuminated the long, axon-like (Fig. 1C, Supplement Video 1), or multiple shorter basal cell processes of MD cells (Fig. S1A). Importantly, truly single-cell Ca^2+^ imaging revealed the presence of subcellular, micro-domain Ca^2+^ transients in both the MD cell somata and processes preceding the global Ca^2+^ spike (Supplemental Video 1). A few MD cells showed pacemaker-like regular Ca^2+^ oscillations with the plateau showing ~4-fold elevations in baseline Ca^2+^ and an average frequency of 0.03/s (Fig. 1C). The average duration of single Ca^2+^spikes (full width at half maximum) was ~2s.

Tissue volume rendering in 3D was performed from z-sections of optically cleared whole-mount MD-GFP kidneys (using the related MD-GFP mouse model that was established recently (Riquier-Brison et al., 2018)) immunolabeled for endogenous MD-specific GFP expression, and tyrosin-hydroxylase (TH) or calcitonin gene-related peptide (CGRP), markers of sympathetic or sensory nerves, respectively (Fig. 2A). There was close anatomical association between MD cell basal processes and the sympathetic and sensory nerve endings (Fig. 2A). In addition, the local expression of synaptophysin, a major synaptic vesicle protein specifically in MD cells and in nearby nerve endings (Fig. 2A) suggest that MD cells synapse with each other and with the sympathetic and sensory nerves. Consistent with MD synaptic function, acute iv administration of several classic neurotransmitters in MD-GT mice, including the ß isoproterenol, the neuroexcitatory glutamate and neuroinhibitory GABA caused immediate, significant alterations in MD cell Ca^2+^ signaling (Fig. 2B). In addition, MD cells responded to several diverse stimuli with altered steady-state Ca^2+^ and/or firing frequency, including mechanical strain (tubule flow), altered tubular fluid composition (low salt diet), local autacoids (angiotensin II) and systemic neurohormone administration (arginine-vasopressin (AVP), gastrin), and metabolic states (diabetic hyperglycemia)(Fig. 2B).

**Figure 2.**
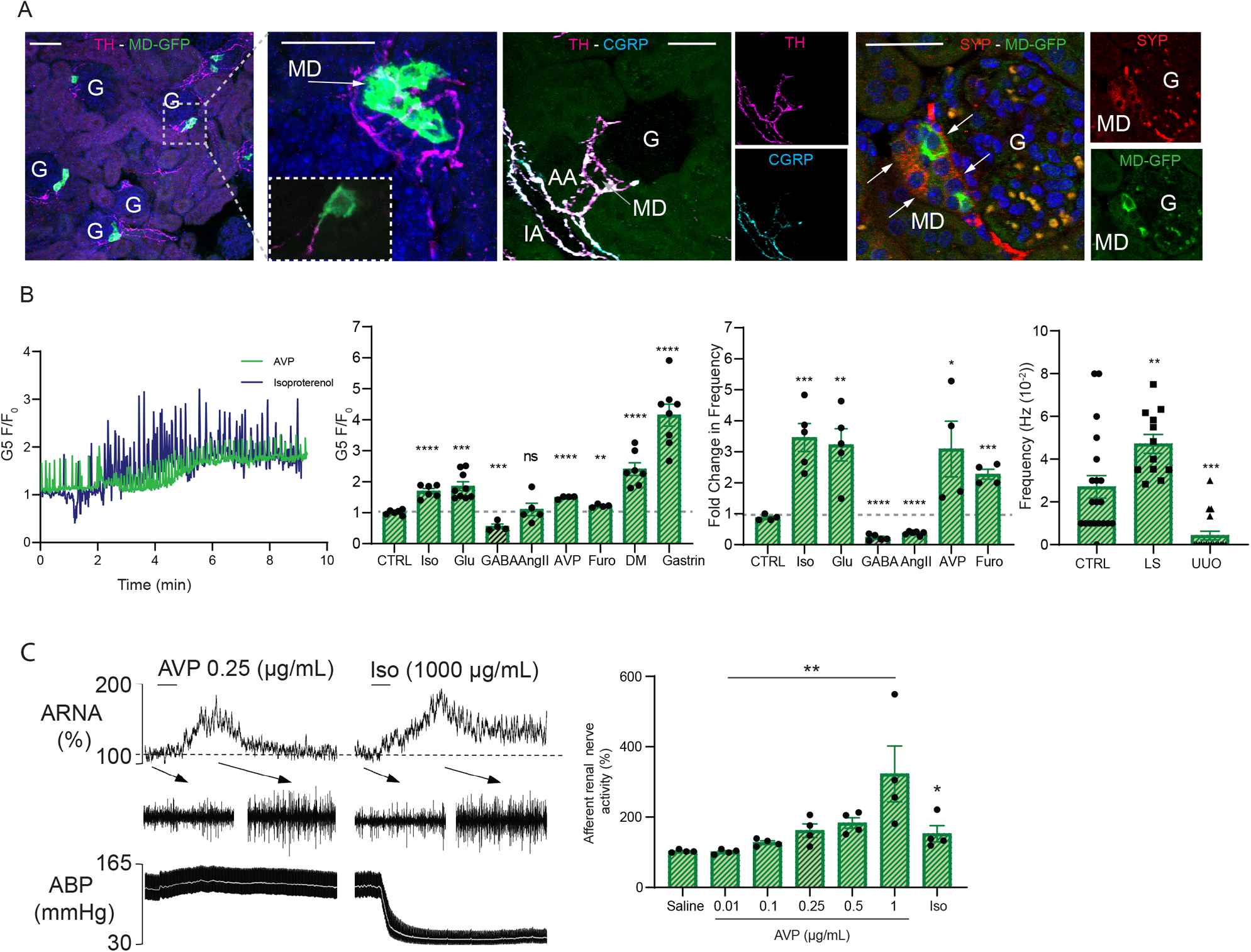
Structural and functional features of MD cell innervation and interoception. **(A)** Tissue 3D volume projection images from optically cleared whole-mount MD-GFP kidneys immunolabeled for endogenous MD-specific GFP expression (green). Left: Tyrosin-hydroxylase (TH) co-labeling (magenta) identifies sympathetic nerve terminals. Magnified area as shown illustrates the close anatomical contact between sympathetic nerve endings and the tip of MD cell basal processes. Center: Co-labeling for TH (magenta) and calcitonin gene-related peptide (CGRP, cyan) that illuminates renal sensory nerves. Overlay and individual TH and CGRP channels are shown separately. Right: Co-labeling for synaptophysin (SYP, red) illuminates the MD (arrows) and the adjacent renal nerve endings (intense red areas). Overlay and individual SYP and MD-GFP channels are shown separately. Bars are 50 μm. **(B)** Multi-cell 4D physiology (mc4DP) mode using Sox2-GT mice. Left: Representativeintensity-based pseudocolor MPI image of a horizontal optical section of a whole-MD (shown in Supplement Video 1). Note the heterogeneity of GCaMP5 fluorescence between individual MD cells. Center: Functional cell-to-cell connectivity map of the same MD plaque and horizontal plane showing all 23 individually numbered MD cell positions. Red line connecting individual cell pairs indicates that the strength of the cell pair correlation (Pearson’s R value) was >0.35. Red/blue cell color indicates hub/lone cells, respectively. Heat-map shows Pearson’s coefficient of each cell pair in a two-color gradient (low correlation blue, high correlation red). Right: Representative simultaneous recording of changes in whole-MD GCaMP5 signal and the adjacent AA diameter over time. **(C)** Afferent renal nerve activity (ARNA) during MD stimulation. Representative traces (left) and statistical summary (right) of arginine-vasopressin (AVP) or isoproterenol (Iso) induced changes in ARNA. Rectified/integrated (0.5 time constant)(top) and raw (1 s examples at baseline and peak response indicated by arrows) (middle) ARNA and arterial blood pressure (ABP) and mean arterial blood pressure (grey line)(bottom). Ns: not significant, *: p<0.05, **p<0.01, data are mean ± SEM, n=4.

We next tested whether MD cells send interoceptive signals to the brain via renal sensory afferent nerves. Intrarenal artery infusion of AVP produced a dose-dependent increase in afferent renal nerve activity (Fig. 2C) and a small increase in arterial blood pressure (saline: −1±1; 0.01ug/mL: 2±1; 0.1ug/mL: 7±2; 0.25ug/mL: 7±3; 0.5ug/mL: 10±3; 1.0ug/mL: 23±13). IV administration of AVP (0.25ug/mL) did not significantly alter afferent renal nerve activity (108±3%, n=3) but produced a significant increase in arterial blood pressure (40±6mmHg, n=3). As a second stimulus, intrarenal artery infusion of isoproterenol also significantly increased afferent renal nerve activity (Fig. 2C) but decreased arterial blood pressure (−73±8 mmHg). To test whether the decrease in arterial blood pressure increased afferent renal nerve activity, IV injection of sodium nitroprusside (100ug/mL, 50uL) significantly decreased arterial blood pressure (−50±3mmHg) but did not alter afferent renal nerve activity (100±2%, n=3).

### Neuron-specific gene profile of MD cells

We next aimed to identify the molecular signature of the neuron-like differentiation and function of the MD. Approximately 28,000 MD cells (representing a very minor fraction, ~0.2% of the total kidney cortical cell population) and 50,000 control cells from adjacent tubule segments were isolated from the cortex of freshly digested MD-GFP kidneys (n=4) for bulk RNA isolation, sequencing and transcriptome analysis (Fig. 3A). Characterization and validation of the bulk RNA sequencing approach, including the high-level expression of known MD cell markers are shown in Figs. 3A and S2. To perform unbiased tissue specificity analysis of the MD cell gene profile (available in NCBI GEO database with GEO accession number GSE163576), we applied TissueEnrich, a recently developed online available tissue-specific gene enrichment analytic tool (Jain and Tuteja, 2019). Using the top 50 highest enriched MD-specific genes as an input gene set, TissueEnrich assigned a highly significant brain tissue identity to MD cells (Fig. 3B). The enrichment of MD-specific genes was highest in the cortex, cerebellum, developing brain, and the olfactory bulb, while kidney-specificity was ranked at only 4 among all organs studied (Fig. 3B). As an alternative method, analysis by the GTEx Multi Gene Query platform of same MD gene set confirmed brain tissue specificity of the MD gene profile (Fig. 3B). In contrast, TissueEnrich analysis of the transcriptome of control kidney cells showed high ranking of kidney specificity and no expression in brain (Fig. S2C).

**Figure 3.**
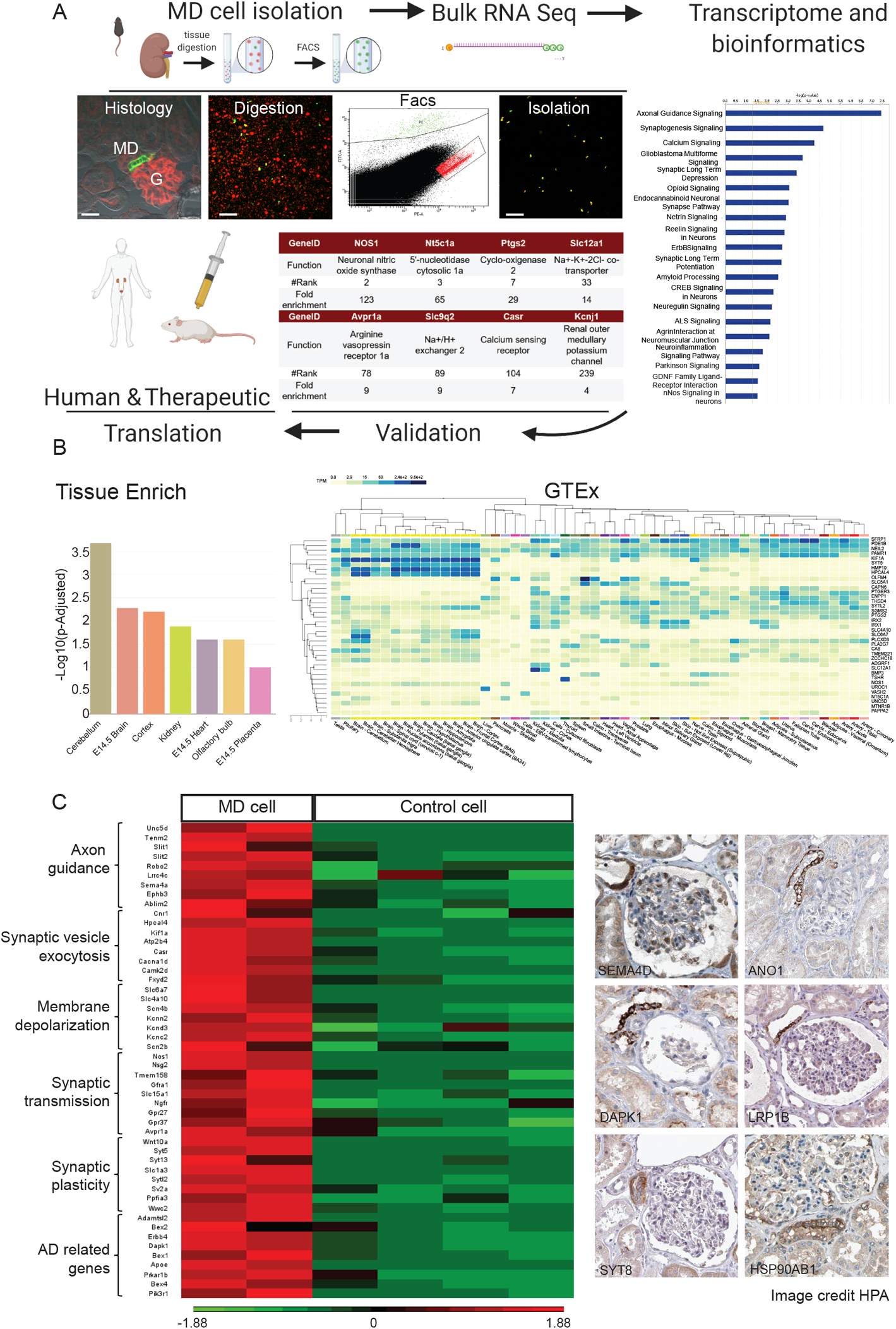
Neuron-like features of the MD cell gene profile. **(A)** Workflow of the MD cell transcriptome analysis. MD (green) and control (red) cells were freshly harvested and digested into single cell suspension from MD-GFP mouse kidneys, and cells were analyzed and sorted based on their fluorescence protein expression (GFP for MD, tdTomato for control cells) followed by bulk RNA isolation and sequencing. Transcriptome analysis showing the most significant MD-specific canonical pathways as listed (based on IPA), and validation of MD specificity by expression of known MD genes (gene names, function, ranking and fold enrichment shown). The expression of specific MD genes was translated to human and therapeutic conditions. **(B)** Tissue specificity analysis of the MD transcriptome. Left: Bar chart showing the enrichment (−Log10(P-Adjusted)) of the top 50 highest expressed MD cell-specific genes in various tissues using TissueEnrich and the Mouse ENCODE dataset for comparison. Right: Heat map of the expression of the same top MD cell gene set in various tissues using GTEx Multi Gene Query. **(C)** The expression of neuronal genes in MD cells. Left: Heat map of the MD (red) vs. control renal cell (green) expression of the top 50 neuron-specific MD cell enriched genes in 6 GO term categories as indicated based on Partek Flow analysis (n=2 MD and n=4 control). **(D)** Immunohistochemistry validation of the human kidney expression and MD cell specificity of top neuronal genes that were identified in the mouse MD transcriptome, including Semaphorin 4D (SEMA4D), Anoctamin (ANO1), Death-associated protein kinase (DAPK1), LDL-related peptide 1B (LRP1B), Synaptotagmin 8 (SYT8), Heat shock protein 90 alpha family class B member 1 (HSP90AB1). Data from the Human Protein Atlas.

To gain detailed molecular insights of the neuron-like MD gene profile, we first performed canonical pathway analysis using IPA. Axon guidance was the most significant pathway, with the high ranking of several other neuron-specific mechanisms (Fig. 3A). Using Gene Ontology (GO) terms that best describe biological processes in neurons as suggested by MD Gene Set Enrichment analysis (Partek Flow), we then selected 7-9 from the top 300 MD-enriched genes into each of the following categories: axon guidance (GO: 0008040), induction of synaptic vesicle exocytosis by positive regulation of presynaptic cytosolic calcium ion concentration (GO: 0099703), membrane depolarization (GO: 0051899), positive regulation of synaptic transmission (GO: 0050806), regulation of neuronal synaptic plasticity (GO: 0048168), and several genes related to neurodegenerative diseases including Alzheimer’s disease (AD). A heat map of these top 50 neuron-specific enriched genes was generated to illustrate their higher and MD-specific expression in MD vs control renal cells (Fig. 3C). The top up-regulated neuron-specific MD genes included previously established MD cell markers (*Nos1, CaSR, Atp2b4, Fxyd2*) as well as new ones (*Unc5d, Tenm2, Hpcal4, Slc6a7, Nsg2, Ngfr, Syt5, Bex1, Dapk1, Apoe*). The highly MD-specific expression of a selection of these genes was validated and translated to the human kidney on the protein level (Fig. 3C).

We next performed single-cell RNA sequencing and transcriptome analysis of individual MD cells harvested from a single MD-GFP mouse to gain additional molecular-level knowledge of the heterogeneity of MD cell biology (GEO accession number GSE189954 for transcriptome data from n=4 different mice). After performing quality control of the initial database, 17,000 genes were detected across 894 MD cells. Unsupervised graph-based clustering and UMAP visualization using Partek Flow revealed five subtypes of MD cells (MD1-5, Fig. 4A). These MD clusters were annotated using a hybrid approach that combined the top enriched genes for each cluster (Fig. 4B-D), classic MD markers (e.g. *Nos1*), cluster-specific canonical pathways based on IPA analysis of the differentially expressed genes (Fig. 4C), and their relevance to the functional sc4DP-based Ca^2+^ signaling results (Fig. 1). The MD1 (high *Itpr1* and *Nfatc2*) and MD3 (high *Pvalb*, *Calb1* and *Grin2c*, glutamatergic signaling) clusters (Fig. 4A-D) likely represent the high Ca^2+^ signaling activity hub and leader (pacemaker) cells that were identified in the in vivo sc4DP functional approach (Fig. 1B). The MD2 cluster (high *Gabrr2* and *Begain*) features high GABAergic signaling and synaptic activity and transmission (Fig. 4A-D) that matches the in vivo functional activity of non-connected “lone” cells (Fig. 1B). The MD4 cluster (high *Apoe*) features the highest expression of genes (*Dscam*, *Axdnd1*) that participate in neuronal cell processes formation, arborization and neurite outgrowth that is consistent with a subset of MD cells having axon-like long basal cell processes (maculapodia) (Gyarmati et al., 2021). The MD5 cluster (high *Nos1*) has the highest expression of the classic MD cell marker *Nos1* and may participate in cell motility, phenotypic transformation and inflammation (Fig. 4A-D).

**Figure 4.**
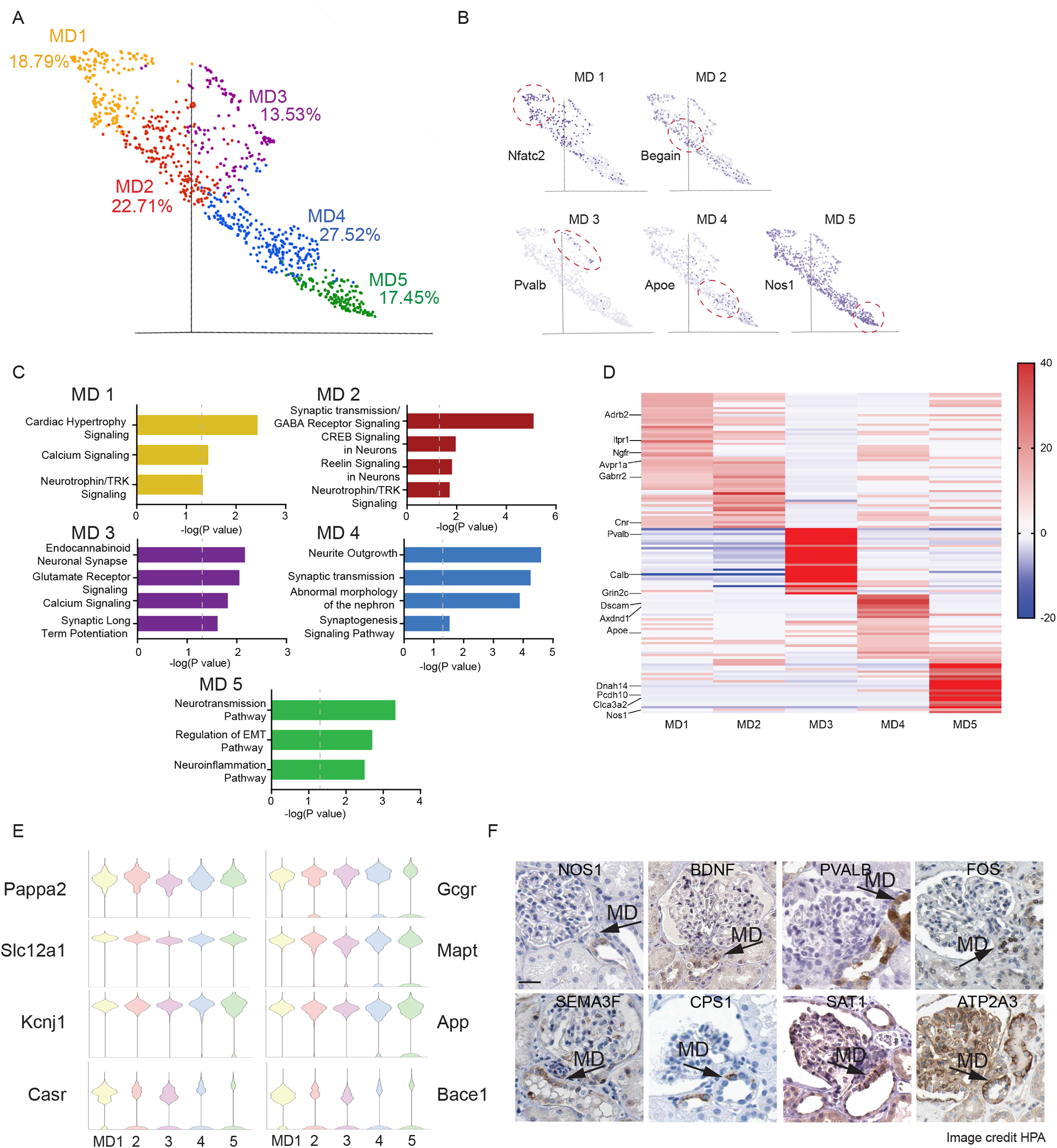
Single-cell transcriptome and heterogeneity of the MD. **(A)** UMAP visualization of integrated single-cell transcriptomic analysis of MD cells (n = 2500). Five cell clusters (MD1-5) were identified based on graph-based analysis in Partek Flow. **(B)** Cluster-specific expression of top enriched identifier genes. These include Nuclear Factor Of Activated T Cells 2 (Nfatc2), Brain Enriched Guanylate Kinase Associated (Begain), Parvalbumin (Pvalb), Apolipoprotein E (Apoe), Nitric Oxide Synthase 1 (Nos1). **(C)** Canonical pathway analysis (based on IPA) showing enriched biological processes specific for clusters MD1-5. **(D)** Heat map of the top 25 enriched genes in each MD cluster (MD1-5). Cluster-specific genes include Adrenoceptor Beta 2 (Adrb2), Inositol 1,4,5-Trisphosphate Receptor Type 1 (Itpr1), Nerve Growth Factor Receptor (Ngfr), Arginine Vasopressin Receptor 1A (Avpr1a), Gamma-Aminobutyric Acid Type A Receptor Subunit Rho2 (Gabrr2), Cannabinoid Receptor 1 (Cnr), Parvalbumin (Pvalb), Calbindin 1 (Calb), Glutamate Ionotropic Receptor NMDA Type Subunit 2C (Grin2c), DS Cell Adhesion Molecule (Dscam), Axonemal Dynein Light Chain Domain Containing 1 (Axdnd1), Apolipoprotein E (Apoe), Dynein Axonemal Heavy Chain 14 (Dnah14), Protocadherin 10 (Pcdh10), Chloride Channel Accessory 3A2 (Clca3a2), Nitric Oxide Synthase 1 (Nos1). **(E)** Violin plot of the expression of highly enriched MD genes in all five clusters (MD1-5). **(F)** Immunohistochemistry validation of the human kidney expression and MD cell specificity of exemplary neuronal genes that show highly heterogenous MD cell expression. These include Nitric Oxide Synthase 1 (NOS1), Brain Derived Neurotrophic Factor (BDNF), Parvalbumin (PVALB), Fos Proto-Oncogene, AP-1 Transcription Factor Subunit (FOS), Semaphorin 3F (SEMA3F), Carbamoyl-Phosphate Synthase 1 (CPS1), Spermidine/Spermine N1-Acetyltransferase 1 (SAT1), ATPase Sarcoplasmic/Endoplasmic Reticulum Ca2+ Transporting (ATP2A3). Data from the Human Protein Atlas.

All MD cells showed high expression of classic MD cell markers (*Nos1*, *Pappa2*, *Slc12a1*, *Kcnj1*, *Casr,* Figs. 4B-E), the newly identified glucagon receptor (*Gcgr*), and a neuron-like gene profile including some of the major AD risk genes (*App*, *Mapt*, *Bace1, Apoe,* Fig. 4D-E). The heterogenous, highly MD-specific expression of a selection of these genes was validated and translated to the human kidney on the protein level (Fig. 4F). The high *NOS1*^+^ cells are consistent with cluster MD5, *BDNF*^+^ cells with MD1-2, *PVALB*^+^ and *ATP2A3*^+^ cells with MD3, *SEMA3F*^+^ with MD4 (Fig. 4F). *CPS1*^+^ and *SAT1*^+^ cells that may synthesize GABA from putrescine via an alternative enzymatic pathway (Kim et al., 2015) may be consistent with MD2 (Fig. 4F). The high level and MD specificity of *FOS* expression is consistent with neuron-like activity, because it is often expressed when neurons fire action potentials (Dragunow and Faull, 1989).

### Nerve Growth Factor (Ngf) regulation of MD cells and kidney function

The nerve growth factor receptor (*Ngfr*) was the highest expressed growth factor receptor in MD cells (ranked #77 most enriched MD gene) suggesting the primary role of Ngf signaling via Ngfr in the neuronal differentiation and function of MD cells. Therefore, we next aimed to characterize the expression, signaling, and function of MD Ngfr. First, a new immortalized mouse MD cell line that we named mMD^geo^ was generated for cell biology studies *in vitro* by following a commonly used workflow as illustrated in Fig. 5A. In addition, we developed mice with MD-specific knockout of Ngfr (MD-NGFR KO) to examine the functional role of MD cell Ngfr signaling *in vivo*. The mMD^geo^ cell line development consisted of primary culturing of freshly isolated and sorted MD cells from MD-GFP kidneys, followed by transfection with LentiSV40 tsA58 for temperature-sensitive proliferation (at 33 ° C for two weeks) in the presence of Ngf. As with the parent MD-GFP mouse model, the cultured mMD^geo^ cell monolayer retained membrane-targeted GFP expression and had a cobblestone pattern typical for epithelial cells (Fig. 5A-B). The MD phenotype of mMD^geo^ cells was validated by the high expression of classic MD cell markers including Cox2, Nkcc2, and Romk (Fig. S3A-B). Differentiated mMD^geo^ cells had almost 10-fold higher expression of Cox2 and Nkcc2 compared to either undifferentiated mMD^geo^ cells grown at 33°C or the formerly established but now extinct MD cell line MMDD1 (Yang et al., 2000)(Fig. S3B). As validation of their intact physiological function, mMD^geo^ cells exhibited the classic phenotype of increased pERK1/2, Cox2, and pp38 expression in response to low salt culture conditions (Fig. S3B). Another classic hallmark of MD cells, Nos1-mediated NO release, and Cox2-mediated PGE_2_ production was confirmed intact in mMD^geo^ cells (Fig. S3C). Interestingly, the heterogeneity of the MD cell population was preserved in the MD^geo^ cell culture with cells showing variable size and shape (Figs. 5A-B, S3A-C), neurotrophin receptor, Cox2 and Nkcc2 expression (Figs. 5, S3A), and NOS activity (Fig. S3C). Supplementation of the MD^geo^ culture media with NGF significantly increased cell proliferation (Fig. 5C) and was absolutely essential for the long-term survival of MD^geo^ cells in culture (not shown).

**Figure 5.**
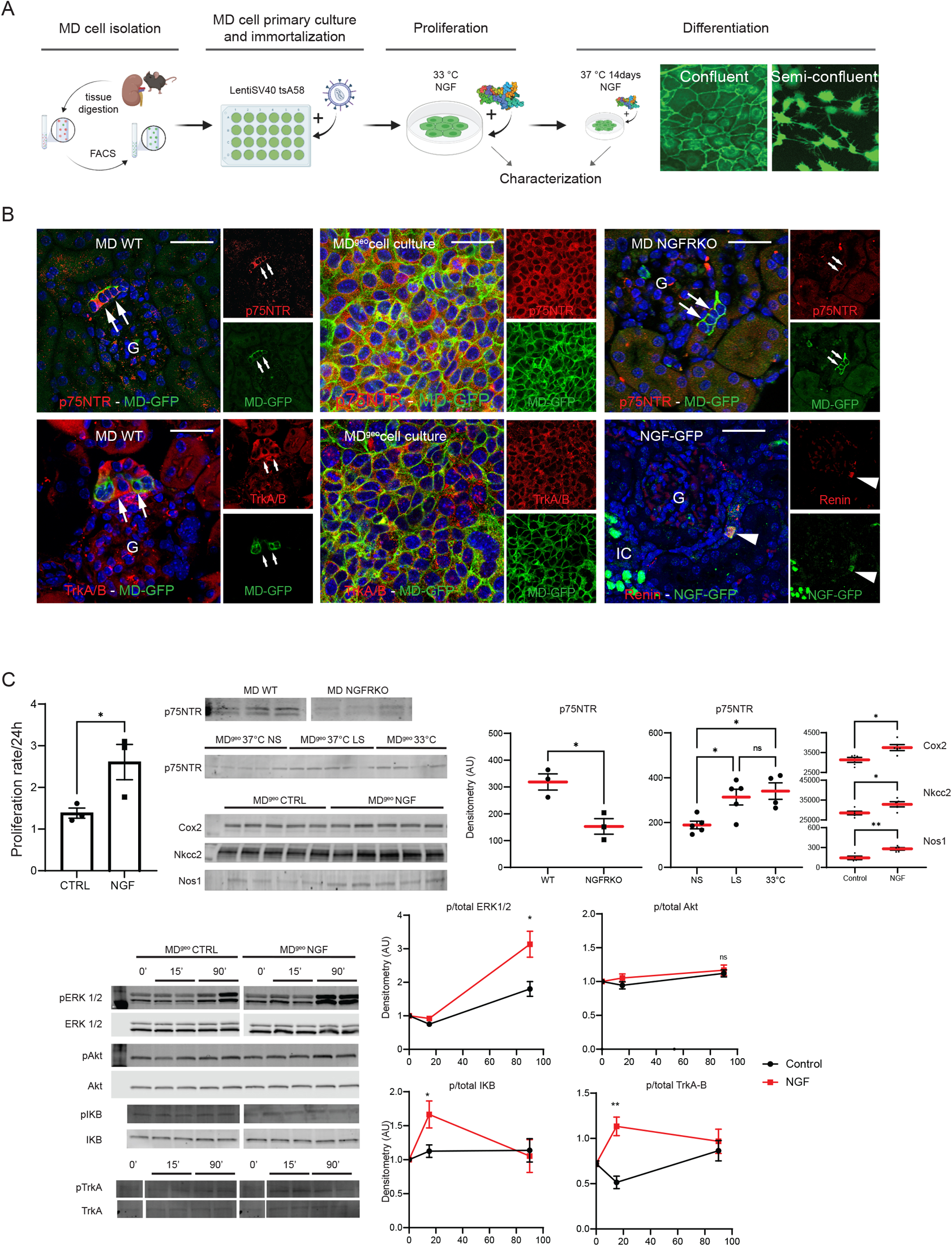
The expression, signaling, and function of Ngfr in MD cells. **(A)** Generation of the novel immortalized mouse MD cell line mMD^geo^. Left: In the workflow, MD-GFP mouse kidneys were freshly harvested and digested into single cell suspension, and cells were analyzed and sorted based on their fluorescence protein expression (MD cells green, all other kidney cell types red). MD primary cultures were immortalized using LentiSV40 tsA58, and either proliferated at 33°C or placed at 37°C for 14 days for differentiation in the presence of NGF followed by characterization. Right: Representative images of fully differentiated mMD^geo^ cells showing epithelial cobblestone-like pattern, with their cell membrane labeled green with endogenous genetic mGFP expression. Semi-confluent MD^geo^ cells feature long, axon-like cell processes. **(B)** Representative immunofluorescence images (overlay, green, and red channels) of the p75NTR Ngfr (red, top row) showing MD cell-specific labeling (arrows, MD identified by genetic GFP expression (green)) in WT mouse kidney (left), in mMD^geo^ cells (center) and in MD-NGFR KO mouse kidney (right). Note the co-labeling of membrane-targeted GFP and Ngfr in both native MD cells in WT kidney and MD^geo^ cells (yellow in overlay) indicating Ngfr membrane localization, but the lack of Ngfr labeling in the MD-NGFR KO mouse kidney. Immunofluorescence images of the TrkA/B Ngfr expression in MD cells (arrows, bottom row) in WT mouse kidney (left), in mMD^geo^ cells (center) and localization of NGF expression in renin-producing JG cells (arrowhead) in the kidney of NGF-GFP reporter mice (right). Native GFP fluorescence is shown in NGF-GFP reporter mouse kidney section, while renin immunofluorescence identified JG renin cells (red). Note the additional strong NGF-GFP signal in intercalated cells (IC) of the renal collecting duct. G: glomerulus. Cell nuclei are labeled with DAPI (blue). Bars are 50 μm. **(C)** The effects of NGF on MD cell function and signaling. Top left: The effects of NGF treatment (0.1 μg/mL) on the proliferation of mMD^geo^ cells. Center/right: Representative immunoblots and statistical summary of p75NTR Ngfr receptor expression in WT and MD-NGFR KO mouse kidney cortex, and in normal salt (NS) and low-salt (LS) treated mMD^geo^ cells cultured at 37°C or 33, and Cox2, Nkcc2, and Nos1 expression in control and NGF-treated mMD^geo 37°C^ cells using whole cell lysates. Bottom: Time-dependent effect of NGF treatment (0.1 μg/mL) on MD cell MAPK, Akt, NFkB and TrkA signaling using mMD^geo^ cells and immunoblotting. Baseline (0’), 15 (15’) and 90 minute (90’) duplicates are shown for phospho (p) or total ERK1/2, Akt, IKB, and TrkA. Statistical summaries show p/total ratio for ERK1/2, Akt, IKB, and TrkA. Ns: not significant, *: p<0.05, **: p<0.01, data are mean ± SEM, n=4-5 each.

Immunofluorescence analysis confirmed MD cell-specific expression of the p75NTR Ngfr in the mouse kidney and in mMD^geo^ cells, including in the cell membrane (Fig. 5B). Both immunofluorescence localization and renal cortical tissue immunoblotting validated the successful MD-specific deletion of the p75NTR Ngfr in MD-NGFR KO mice (Fig. 5B-C). Compared to fully differentiated mMD^geo^ cells, Ngfr expression was significantly increased by low salt culturing condition and in rapidly proliferating undifferentiated mMD^geo^ cells (Fig. 5C). Importantly, NGF treatment significantly increased the expression of the classic and functionally key MD cell markers Cox2, Nkcc2, and Nos1 (Fig. 5C). NGF treatment induced phosphorylation of ERK1/2 MAP kinases, IKB and TrkA-B, but not Akt in a variable time-dependent manner (Fig. 5C). Using NGF-GFP reporter mice, the expression of the Ngfr ligand NGF in renin producing JG cells was confirmed (Fig. 5B). Additional strong NGF expression was found in intercalated cells of the renal collecting duct (Fig. 5B).

MD cells are classic key controllers of basic kidney functions such as glomerular filtration rate (GFR) and renin secretion (activation of the renin-angiotensin system). Therefore, we finally tested the functional role of MD Ngfr using MD-NGFR KO mice and the effect of exogenous NGF treatment *in vivo*. Compared to WT mice, the frequency of MD cell Ca^2+^ transients showed a 4-fold increase in MD-NGFR KO mice (Fig. 6A). Importantly, the regular periodic oscillations in whole-MD Ca^2+^ that were observed in WT mice (Fig. 1B) were replaced by irregular chaotic oscillations in MD-NGFR KO mice (Fig. 6B). Cell-to-cell connectivity of MD cells was greatly reduced in MD-NGFR KO mice compared to WT (Fig. 1B) as shown by the connectivity map and heat map that were generated based on Pearson’s correlation analysis of the time-lapse Ca^2+^ recordings between all cell pairs (Fig. 6B). In the MD of MD-NGFR KO kidney, there were only two small “hub” cells within the whole MD, while a great number of “lone” cells were identified which had no connections to other MD cells (Fig. 6B). In addition, MD cells in MD-NGFR KO mice generated significantly reduced Ca^2+^ responses to iv injected gastrin (Fig. 6B). The injection of NGF, or the p75NTR receptor modulator LM11A-31, or recombinant soluble β-amyloid precursor protein alpha (sAPPα) all induced significant elevations in MD cell Ca^2+^ (but not in any other renal cell types, not shown) in the WT but not in MD-NGFR KO mice (Fig. 6C). The exception was NGF which produced strong MD cell Ca^2+^ responses not only in WT, but also in MD-NGFR KO mice (Fig. 6C). Interestingly, the expression of APP (precursor of sAPPα) (Fig. 6C) in addition to the other Ngfr ligand NGF (Fig. 5B) was confirmed in renin producing JG cells. Short-term treatment with NGF (10 µ kg sc daily for one week) caused significant increases in the number of renin-producing juxtaglomerular (JG) cells and in GFR in WT mice, while renin cell number and GFR were significantly lower in MD-NGFR KO mice (Fig. 6D). In addition, MD-NGFR KO mice featured kidney tissue fibrosis and showed increased levels of p-tau (S199) in MD cells (Fig. 6E).

**Figure 6.**
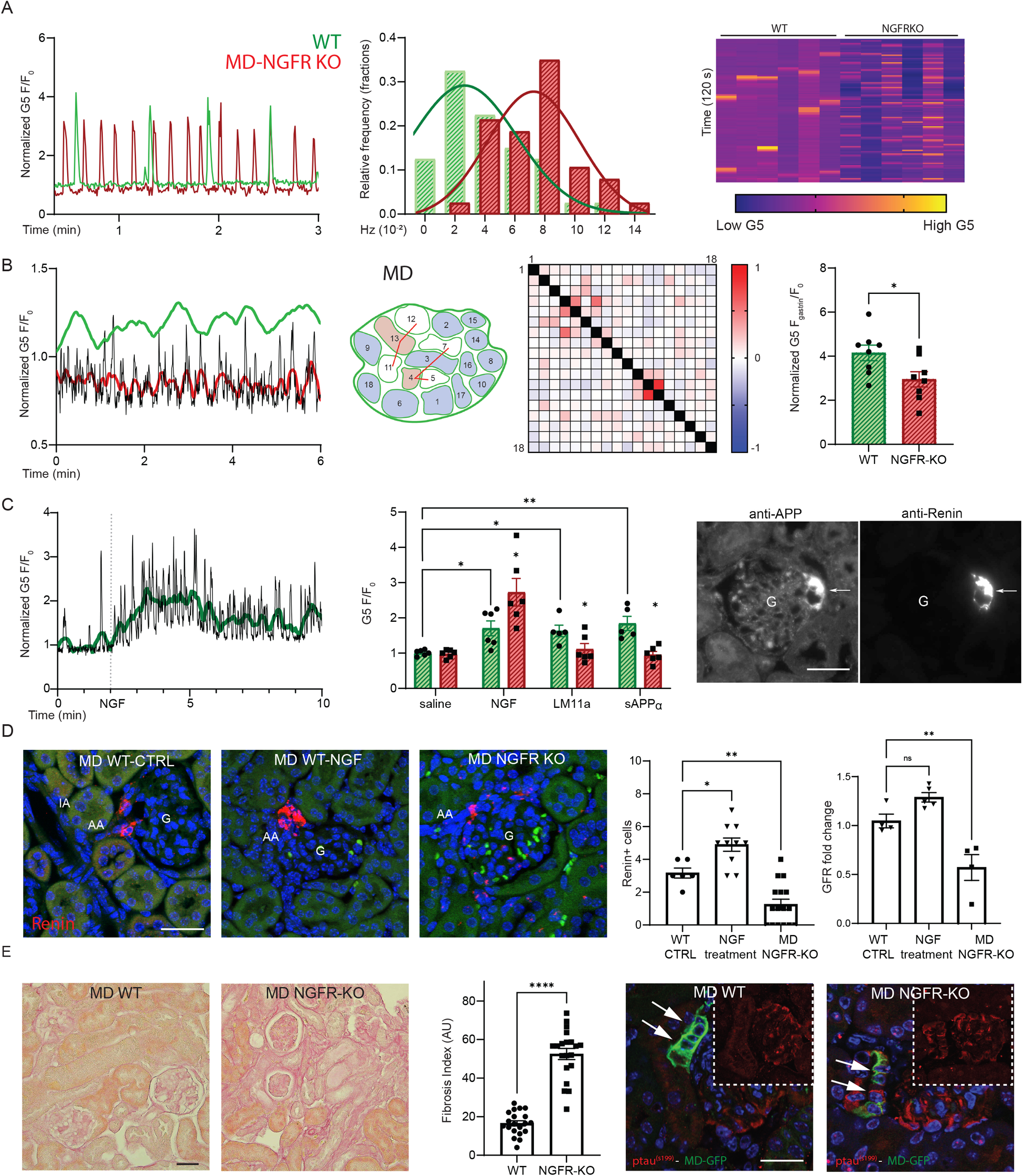
The function of MD cell Ngfr *in vivo*. **(A)** Increased frequency of MD Ca^2+^ transients in MD-NGFR KO vs. WT mice. Left: Representative recordings of the regularly oscillating Ca^2+^ transients in a single MD cell from WT (green) and MD-NGFR KO mouse (red). Center: Distribution of single MD cells (shown as fraction of total number of cells with curve fittings) based on their Ca^2+^ firing frequency during 2-min time lapse recordings. Right: Heat map of the changes in GCaMP5 fluorescence intensity (F/F_0_) of six individual MD cells from both WT and MD-NGFR KO mice during a 120 s time-lapse. Note the 4-fold increase in Ca^2+^ firing frequency in MD-NGFR KO vs WT. **(B)** Reduced connectivity and sensitivity of MD cells in MD-NGFR KO vs WT mice. Left: Representative recordings of changes in whole-MD GCaMP5 signal over time in a WT (green) and a MD-NGFR KO mouse (red). Center: Functional cell-to-cell connectivity map of a MD plaque shown in horizontal plane and with all 19 individually numbered MD cell positions. Red line connecting individual cell pairs indicates that the strength of the cell pair correlation (Pearson’s R value) was >0.35. Red/blue cell color indicates hub/lone cells, respectively. Heat-map shows Pearson’s coefficient of each cell pair in a two-color gradient (low correlation blue, high correlation red). Right: The effect of iv injected gastrin (5 ug/mL) on MD cell Ca^2+^ (G5 F/F_0_) in WT (green) vs. MD-NGFR KO mice (red). *: p<0.05, data are mean ± SEM, two data points from n=4 mice each. **(C)** The effects of iv injected exogenous Ngfr ligands on MD cell Ca^2+^ in WT (green) and MD-NGFR KO mice (red). Left: Representative time-lapse recording of the effect of NGF (0.25 ug/g) on the whole-MD calcium (G5 F/F_0_) in a WT mouse. Center: Summary of the effects of iv injected NGF (0.25 ug/g), or LM11A-31 (75 ug/g), or sAPPα (0.6 ug/g) on fold-changes in MD cell Ca^2+^ (G5 F_max_/F_0_) in WT (green) vs. MD-NGFR KO mice (red). *: p<0.05, **:p<0.01, data are mean ± SEM, n=5 mice each. Right: Immunofluorescence images of APP and renin double-labeling in renin-expressing JG cells (arrow) in a mouse kidney section. G: glomerulus. (D) Reduced renin expression and GFR in MD-NGFR KO vs WT mice. Left: Representative immunofluorescence images and summary of the effects of NGF treatment for one week (0.25 ug/g/day) on renin (red)-producing JG cell number in WT untreated, WT NGF treated, and MD-NGFR KO mice. Cell nuclei are labeled blue with DAPI, tissue autofluorescence (green) is shown for tissue morphology. Right: Summary of the changes in GFR in untreated WT, WT treated with NGF, and MD-NFGR KO mice. *: p<0.05, **: p<0.01, data are mean ± SEM, one average shown from n=6-10 mice each. **(D)** Tissue fibrotic and MD neurodegenerative features of MD-NGFR KO mice. Left: Representative images of PAS-stained kidney tissue sections and the summary of tissue fibrosis index in WT and MD-NGFR KO mice. ****: p<0.0001, data are mean ± SEM, four data points are shown from n=5 mice each. Right: Immunofluorescence images of p-tau expression (red) in MD cells (arrows) of WT and MD-NGFR KO mice. Overlay with endogenous MD-GFP signal (green), red channel shown separately in insets. Cell nuclei are labeled blue with DAPI.

## DISCUSSION

This study uncovered new neuron-like functional and molecular features of MD cells in the kidney and established their fundamental roles in interoception, physiology and disease. Our results show that MD cells are chief sensory neuroepithelial cells that form an autonomous neural network that functions as the nephron central command to control key organ and whole body functions. As interoceptive neuron-like cells in the kidney, MD cells sense the local and systemic environment, process and send signals to renal and central effectors to maintain homeostasis. Several major research technology innovations developed in this work made it possible to (i) directly visualize the unique Ca^2+^signaling and pacemaker activities of single MD cells in vivo (using sc4DP) that are altered by sensing the systemic and local tissue environment via multi-organ crosstalk, (ii) establish the neuron-like gene profile of MD cells (using bulk and scRNA sequencing) providing molecular detail of their heterogenous biological and physiological functions, and (iii) identify NGF signaling as the principal regulatory mechanism of MD cells and their control of key kidney functions and role in disease development (using the new MD cell line mMD^geo^ and MD-NGFR KO mice). Due to the molecular-level new knowledge of multi-organ relevant functions of MD cells established here, our new MD cell research toolbox will be useful for future cell biology studies in the kidney, heart, intestine, pancreas and brain and directly relevant to modeling kidney and neurodegenerative diseases.

Intravital imaging of the activity of MD versus all other renal cell types using calcium signaling as a readout established MD cells as the most active, chief cell type in the kidney cortex. The presently uncovered new cell physiology features including the autonomous, rhythmic Ca^2+^ firing (pacemaker function), the coordination and clustering of single MD cell Ca^2+^ transients into a cumulative whole-MD signal that translates to the regulation of key whole-organ functions (e.g. blood flow) further suggest that MD cells act as the nephron central command. MD cells are known to drive the TGF component of the regular physiological oscillations in renal blood flow and glomerular filtration rate that has an established frequency of 0.03 Hz and cycle time of 30-50 s (Marsh et al., 2019). The oscillation frequency of whole-MD Ca^2+^ signal is a perfect match with that of TGF, and the peak MD Ca^2+^ is phase-matched with the most dilated state of the AA (Fig. 1B). These results suggest that one of the many potential functions of MD Ca^2+^ signals is the periodic generation of vasodilator effectors (likely NO and PGE_2_) that cause paracrine AA dilatations, increased blood flow and glomerular filtration. In fact, the Nos1 and Cox2 enzymes that generate these vasodilators are known classic MD-specific markers in the renal cortex (Harris et al., 1994; Mundel et al., 1992; Peti-Peterdi and Harris, 2010) and known to be Ca^2+^ sensitive (Abu-Soud et al., 1994; Zhou et al., 2014). Importantly, MD Ca^2+^ activity increased in response to stimuli that cause low salt in the tubular fluid microenvironment including low salt diet and the diuretic furosemide that inhibit MD NaCl entry via Nkcc2 (Slc12a1) (Fig. 2B). This is consistent with the classic function of MD cells in tubular fluid salt sensing that is known to involve Nkcc2 and MD cell depolarization in response to altered luminal [NaCl] (Lapointe et al., 1990). In addition to tubular salt, MD cells were able to directly sense numerous diverse signals from the local tissue and systemic environment and to process them and modulate their Ca^2+^ activity accordingly, suggesting their major function in interoception. The MD Ca^2+^ agonist/antagonist profile (Fig. 2B) suggests that MD cells can detect variations in sympathetic tone (ß-agonist isoproterenol, Iso), systemic neurohormones that signal altered body fluid volume or osmolality and postprandial metabolic states (AVP, angiotensin II, gastrin), metabolic disease states (diabetic hyperglycemia), and local mechanical strain (tubule flow) and based on these signals MD cells regulate organ blood flow and glomerular filtration. Some of these findings were expected and consistent with the known MD mechanosensory (Sipos et al., 2010) and metabolism sensing functions (Tonneijck et al., 2017; Vargas et al., 2009), and with the MD expression of the relevant neurohormone receptors including the V1a AVP receptor as reported earlier (Aoyagi et al., 2008) and the Adrb2 receptor for Iso (Fig. 4D). Others, for example the most significant MD Ca^2+^ agonist actions of gastrin (Fig. 2B) are exciting new findings that hint on direct crosstalk between the kidney and other distant organs such as the intestine. Gastrin sensing by the MD via *Cckbr* may be a new mechanism in the gut-renal axis and in the physiological postprandial adjustments of glomerular filtration (Bankir et al., 2015; Tonneijck et al., 2017). However, it is likely that these examples represent only a fraction of the sensing capabilities of the MD, since the new MD cell gene profile (discussed below) established the expression of several other metabolic and hormonal receptors (e.g. for glucagon, oxytocin, growth hormone, etc). Importantly, MD-specific stimuli (intra-renal AVP and Iso administration) triggered significant dose-dependent increases in ARNA (Fig. 2C). These results functionally confirm that MD cells synapse with sensory afferent nerve endings and by sending signals to the brain via ARNA, MD cells participate in systemic interoception.

Single-cell expression of GT reporters in the MD was made possible by partial tamoxifen induction of the underlying Cre-lox genetic strategy (Gyarmati et al., 2021). This model allowed truly single-cell Ca^2+^ imaging in the sc4DP mode with subcellular resolution and free of signal contamination from neighboring cells (Fig. S1). Due to the technical advantage provided by this sc4DP approach, it became possible to identify regularly spiking pacemaker MD cells (Fig. 1C). Although periodic cell depolarizations as the origin of Ca^2+^ transients (Ca^2+^ firing) were not measured in this study, the role of MD membrane depolarizations in MD cell physiology is well known (Bell et al., 2003; Lapointe et al., 1990). However, Ca^2+^ imaging of these cells has been notoriously difficult using classic calcium fluorophores due to technical constraints (Komlosi et al., 2008; Peti-Peterdi and Bell, 1999). The MD cell pacemaker activity uncovered in the present study resembles those in the interstitial cells of Cajal (ICC) in the intestine and in pacemaker neural networks of other organs including the intestine, pancreatic islets and the sinoatrial node in the heart (Da Silva Xavier and Rutter, 2020; Drumm et al., 2017; Liang et al., 2021). The average duration of single Ca^2+^spikes (full width at half maximum) was ~2s consistent with the slow waves that are typical of ICCs in the small intestine (Drumm et al., 2017). In addition to these functional characteristics, the expression of all key molecular elements required for pacemaker cell activity were identified in the MD transcriptome (Figs. 3–4). The functional map of the MD (MD connectome) identified highly connected hub (or “leader”) cells and of subordinate (or “follower”) or non-connected (“loner”) cell populations (Fig. 1B). Similar cell hierarchy was described recently for a single pancreatic islet (Salem et al., 2019). Overall, these results showed the strong connection and organization of the individual MD cells into a functional unit (syntitium), a neural network that drives the vasomotion of the glomerular afferent arteriole. This in turn regulates renal blood flow, glomerular filtration, tubular fluid flow and hence the function of the entire nephron.

Both functional, (Fig. 1B), structural (Fig. 2A) and molecular genetic evidence of both pre and postsynaptic marker expression (Fig. 3C) suggested the presence of synapses between individual MD cells and sympathetic and sensory nerve endings. Their potential roles include local cell-to-cell communication and coordination within the whole-MD (formation of the MD neural network), direct regulation of MD cell function by sympathetic efferent nerves and providing sensory input to the central nervous system via afferent renal nerves. The presently established neuron-like sensory and interoception MD function may be the kidney-derived mechanism behind the sympathoexcitatory effect of the diuretic furosemide that has been long known to involve renal sensory receptors and afferent renal nerves (Petersen and DiBona, 1995). The physical shape of the MD plaque, a niche of ~20-25 cells appears to match its function as it resembles a peripheral ganglion (Fig. 1B). In fact, the MD neural networks in each nephron that were identified in the present study may represent a new “renal nervous system” controlling kidney function and sensing the internal milieu of the entire body in a novel peripheral interoceptive neural pathway. Functional interconnections of MD neural networks in each nephron and between the local peripheral MD “ganglia” and the central nervous system via sensory nerves are exciting possibilities for exploration in future work.

The MD transcriptome analysis identified the high level and MD-specific expression of several neurohormone and neurotransmitter receptors including *Adrb2*, *Grin2c*, *Gabrr2*, *Avpr1a*, *Cckbr*, etc (Figs. 3–4). These results confirm the functional expression of these receptors that was first established by using their respective ligands in our in vivo Ca^2+^ imaging sc4DP methodology (Fig. 2B), validating the synergistic functional and molecular approaches applied in the present work. In fact, we emphasize the top value and primary importance of the functional, physiological insight (Figs. 1–2) that initiated this study and was the main driver of the overall study design. The observed neuron-like physiology and the functional heterogeneity between individual MD cells (sc4DP) provided the rationale for performing bulk and then scRNA sequencing and transcriptome analysis, which in turn identified Ngfr as the highest expressed MD growth factor receptor and drove the development of MD^geo^ cells and the MD-Ngfr KO mouse model. Clearly, physiological knowledge and context is essential for the pursuit and interpretation of scRNA sequencing-based transcriptome analysis, which is often overlooked in the rather transcriptome-focused present era of biology research.

Importantly, the neuronal features of MD cell transcriptome were found in nearly all MD cells and were not limited to a select minor cell population. In addition, histological analysis (Fig. 3A and (Gyarmati et al., 2021; Riquier-Brison et al., 2018)) confirmed that all GFP^+^ cells in the renal cortex that were sorted and analyzed in the present study in the bulk and single-cell RNA seq-based transcriptome analysis were confined to the MD in the control MD-GFP mouse kidney. These findings argue against the possibility that our samples were contaminated by a few truly neuronal cell types that were reported recently (Lake et al., 2021). The neuroepithelial character of MD cells was suggested recently by a preliminary scRNA sequencing study that analyzed 24 single MD cells (Chen et al., 2021a). A few other recent studies reported limited MD transcriptome data (the expression of *Nt5c1a* and *Slc5a1*) using a very low number of cells (He et al., 2021; Ransick et al., 2019; Zhang et al., 2019).

Based on the presently established MD cell transcriptome, MD cells share dominant biological properties with glutamatergic neurons, similar to other pacemaker cell types in other organs including the sinoatrial node in the heart (Liang et al., 2021), the interstitial cells of Cajal in the small intestine (Lee et al., 2017), and in the pancreatic islets of Langerhans. In addition, MD cells share functional and molecular similarities with Nos1^+^ glutamatergic neurons in the CNS which play important roles in Alzheimer’s disease (Park et al., 2020). Among the several neuron-specific genes highly expressed in MD cells (Fig. 3C) Nsg2 functions as an AMPAR-binding protein that is required for normal synapse formation and/or maintenance (Chander et al., 2019). Several neuronal genes highly expressed in the MD were directly implicated in Alzheimer’s disease that suggest the activity of neurodegenerative mechanisms in MD cells in addition to their physiological neuron-like function. More detailed classification of MD cell subtypes in the future may consider cell morphological and other cell biology features as applied before for Nos1^+^ neurons (Tricoire and Vitalis, 2012).

The transcriptome data identified Ngfr as the highest expressed growth factor receptor in MD cells, and this new knowledge turned out to be essential for the successful generation of the new MD^geo^ cell line and informed the development of the MD-NGFR KO mouse model. Multiple initial attempts to primary culture the freshly isolated MD cells from MD-GFP mouse kidneys failed until NGF was added to the culture media. In fact, NGF promoted MD^geo^ cell proliferation, increased the expression of several classic MD cell markers, induced ERK1/2 and NFκB signaling (Fig. 5C) that is typical of Ngfr (Lin et al., 2015), enhanced MD cell Ca^2+^ signaling (Fig. 6C), and induced renin cell number and GFR (Fig. 6D). This wide range of stimulatory effects are consistent with the known neurotrophic function of NGF to promote the survival, growth, and differentiation of neuronal cells (Amadoro et al., 2021) and with the notion that Ngfr signaling is the major driver of the neuronal differentiation of MD cells. It should be noted that the possible involvement of pro-NGF signaling in these responses cannot be excluded since the p75NTR Ngfr binds all pro-neurotrophins (Hempstead, 2006). Future work is needed to clarify details of the rather complex (pro)NGF signaling in MD cells. The actions of NGF via the p75NTR Ngfr usually involves Trk receptors, although kinase-independent signaling mechanisms have also been identified (Lin et al., 2015). While MD cells have detectable TrkA-B expression (Fig. 5B), the MD transcriptome did not show enrichment of Ntrk1-2 in MD vs control renal tubular epithelial cells in contrast to the robust MD-specific Ngfr levels.

Interestingly, renin producing JG cells that are the immediate neighbors of MD cells have high expression of NGF and APP (Figs. 5B, 6C), two potential Ngfr ligands, that suggest the possibility of new types of paracrine cell-to-cell signaling between MD and JG renin cells. Another source of NGF in the kidney may be Gli1^+^ renal interstitial myofibroblasts (E et al., 2019), but autocrine neurotrophin signaling between specific MD cell subsets is also a possibility (e.g. via BDNF, Fig. 4F).

Immortalized MD^geo^ cells maintained the expression of membrane-targeted GFP similarly to native MD cells in the MD-GFP kidney (Gyarmati et al., 2021; Riquier-Brison et al., 2018; Shroff et al., 2021), validating their MD cell origin. In addition, MD^geo^ cells highly expressed known MD cell markers such as Nos1, Nkcc2, Cox2, and their expression showed physiological regulation (Fig. 5C). Cultured MD^geo^ cells maintain the physiological heterogeneity of native MD cells in cell shape, size, Cox2 and Nkcc2 expression and NO generation, and feature the recently described axon-like basal cell processes (Gyarmati et al., 2021)(Fig. 5A) further validating the functional and transcriptomic data (Figs. 1–4). The immortalized MD cell line MD^geo^ may be useful to study or model not only MD cells in the kidney, but a variety of other cell types throughout the body, e.g. pacemaker cells that form a neural network.

In addition, the high expression of the AD molecular fingerprint (Grubman et al., 2019; Roussarie et al., 2020) in MD cells (Fig. 3C, 4B-E) suggests MD^geo^ cells may be used for studying cell and molecular mechanisms of neurological and neurodegenerative diseases including Alzheimer’s disease. Interestingly, MD cells have the highest overall rate of protein synthesis among all kidney cell types (Shroff et al., 2021). The high level of Tau (*Mapt*) and amyloid (*App*) expression in MD cells (Figs. 4E and 6E) represents several layers of similarity between MD cells and Nos1^+^ neurons, their functionally relevant role in tubulo-glomerular feedback and neurovascular coupling (organ blood flow control), and disease relevant dysfunctions in kidney fibrosis and neurodegenerative diseases, respectively. Tau in the MD may regulate Nos1-derived nitric oxide (NO) generation in the glutamatergic synapse similarly to Nos1 expressing glutamatergic neurons in the CNS, and hence neurovascular (tubulo-glomerular feedback) function (Park et al., 2020). Excess NO generation by GABAergic Nos1^+^ neurons can also lead to their dysfunction that has been implicated in Alzheimer’s disease development (Choi et al., 2018). The present study established that Nos1^+^ MD cells have both glutamatergic and GABAergic synapses and high expression of numerous molecular elements of a robust actin-cytoskeleton and microtubule-associated proteins including tau. In addition, the acute administration of several p75NTR Ngfr agonists (NGF, LM11A-31, sAPPα) demonstrated functional expression of Ngfr in MD cells *in vivo* (Fig. 6C). The small molecule LM11A-31, a specific Ngfr modulator that is currently in human clinical trial had highly promising results in mouse models of Alzheimer’s disease (Simmons et al., 2014). sAPPα is another known p75NTR agonist with neuroprotective effects (Harris et al., 2020). Interestingly, Ngfr deficiency triggered MD hyperactivity at the single cell level (increased Ca^2+^ firing frequency, Fig. 6A) which may have been due to glutamatergic hyperexcitability or the loss or dysfunction of inhibitory GABAergic MD cells that are well-known key, early pathogenic events in the development of neurodegeneration (De Strooper and Karran, 2016). This, in turn, led to the loss of cell-to-cell connectivity and absence of the periodically oscillating vasomotor (TGF) signal at the whole-MD level (Fig. 6B). As a result of altered MD signaling, GFR declined, renin cell number decreased, and renal tissue fibrosis developed (Fig. 6D-E). The irregular chaotic oscillations in whole-MD Ca^2+^ that were observed in MD-Ngfr KO mice (Fig. 6B) were reminiscent of the previously reported chaotic oscillations in MD-mediated TGF in different models of hypertension (Holstein-Rathlou and Leyssac, 1986; Yip et al., 1991). These results suggest that chaotic behavior of the MD neural network is a common pathogenic feature of the dysregulated renal vascular control and blood flow in hypertension and kidney disease. Again, this is in agreement with the known important role of neurovascular dysfunction in the development of neurodegenerative diseases (Kisler et al., 2017). In addition to the dysregulated vascular control function, the present findings (Fig. 6E) suggested that altered NGF signaling in MD cells results in increased levels of phosphorylated tau which is the tell-tale sign of neurodegeneration (Sexton et al., 2021).

Based on the new results, we put forth a new, single-cell MD model and concept of neurodegenerative disease development. This MD model integrates the currently existing major paradigms of neurodegeneration including neurovascular dysfunction (Kisler et al., 2017), neurotrophic (NGF) hypothesis (Amadoro et al., 2021; Appel, 1981), amyloid and tau pathologies (Hardy and Selkoe, 2002; Sexton et al., 2021) integrated in a single cell type (MD). The simplicity of the MD function and model in the kidney makes it easier to study the otherwise highly complex neurodegenerative mechanisms in the brain. In contrast to our single-cell MD model, the current integrated hypothesis for Alzheimer’s disease concerns a rather long, complex multi-cellular communication and responses between neurons, astrocytes, microglia, and the vasculature (De Strooper and Karran, 2016). Therefore, this new single-cell MD model presents both conceptual and technology advantages for our improved understanding of neurodegeneration and for experimental research. In fact, the present study identified several new molecular and mechanistic targets (Figs. 3–4) and developed new *in vitro* (mMD^geo^ cell line, Figs. 5 and S3) and *in vivo* research tools (MD-NGFR KO mice, Fig. 6) to facilitate future research in both kidney and neurodegenerative diseases.

The overall view emerging from our new results suggest that a pattern in the intensity and complexity of environmental signals at the glomerular vascular pole and in the MD cell responses to these stimuli (physiology) have led to the neuronal differentiation of MD cells and caused heterogeneity in the transcriptome and proteome of individual cells (MD clusters). This in turn created cellular hierarchies (connectome) and established the coordinated functional MD network. Due to its many fundamental sensory, vasomotor, and secretory functions (Fig. 7), the MD neural network is functioning as the nephron central command. MD deficiency of Ngfr, the major determinant of its neuron-like function causes dysfunction in the MD neuronal network and leads to disease development that is reminiscent of neurodegeneration.

**Figure 7.**
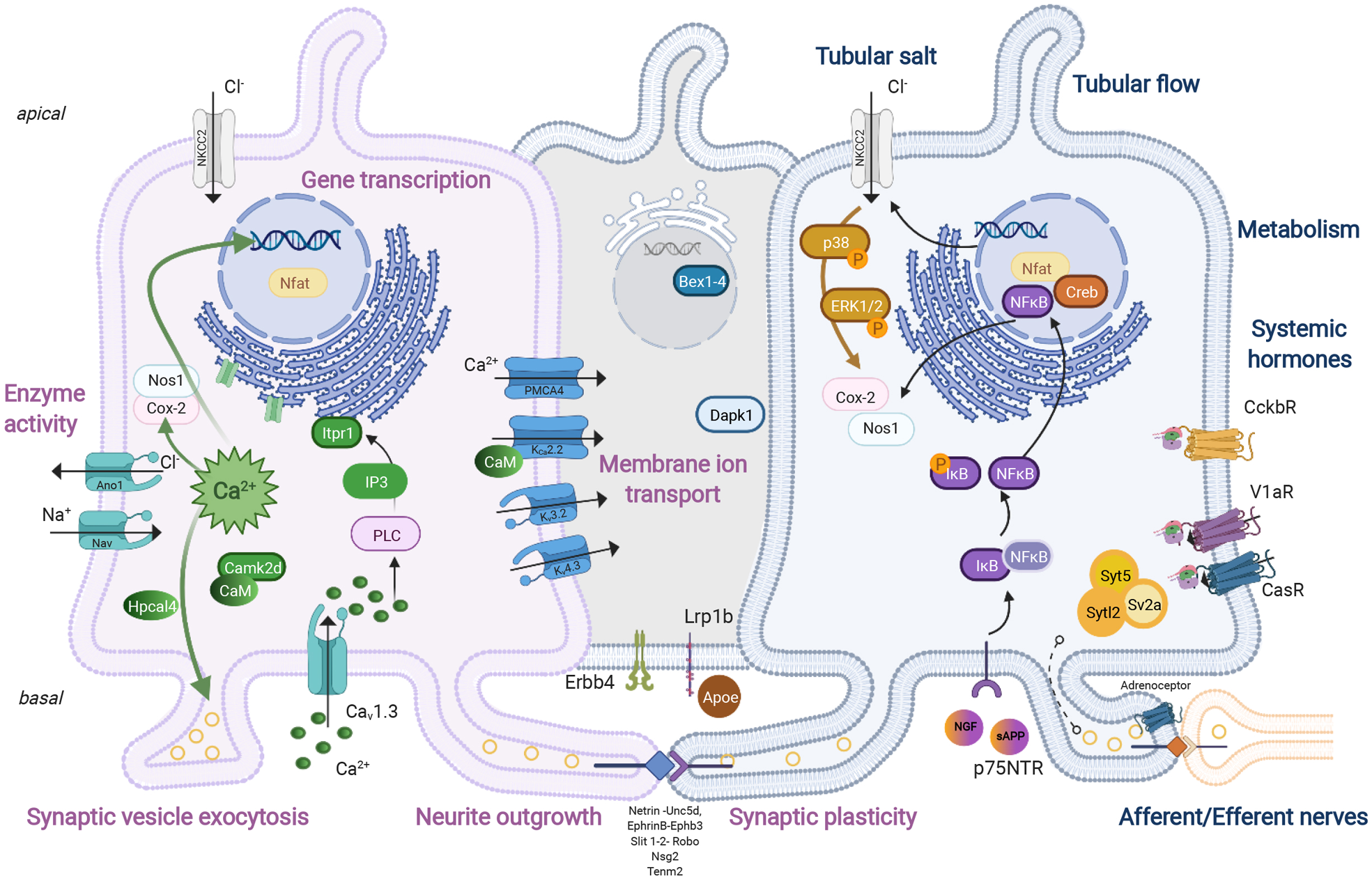
Schematic model of the neuron-like molecular and cell biology of MD cells. The numerous local and systemic environmental factors detected by the sensory MD cells (blue) and the functional role of MD cell calcium and NGFR signaling in cell biology and physiology (purple) are highlighted based on calcium imaging, transcriptome and pathway analysis, in vitro cell culture and in vivo (patho)physiology data acquired in the present study.

## MATERIALS AND METHODS

### Animals

Male and female, 6-8 weeks old C57BL6/J mice (Jackson Laboratory, Bar Harbor, ME) were used in all imaging and histology experiments. Male and female Sprague-Dawley rats (n=2 each, 250-350g, Charles River Laboratories) were used for renal afferent nerve activity measurements and were pair-housed in a temperature-controlled room (22±1°C) with a 12-hour dark:light cycle (lights on at 07:00-19:00) and given ad libitum access to deionized water and 0.1% NaCl chow diet (Research Diets, D17020). Tamoxifen-inducible, conditional macula densa (MD) cell-specific p75NTR knockout (MD-NGFR-KO) mice on a C57BL6/J background were generated by intercrossing nNOS/CreERT2 (Taniguchi et al., 2011) and p75NTR/floxed mice (Bogenmann et al., 2011) (possess *loxP* sites flanking exons 4 through 6 of the *Ngfr* gene, both from Jackson Laboratory). MD-NGFR-KO mice were further backcrossed with GCaMP5-tdTomato (GT)/floxed reporter mice (Gee et al., 2014) resulting in the expression of the calcium sensitive green fluorescence protein GCaMP5 and the calcium insensitive fluorescence protein tdTomato specifically in MD cells (Jackson Laboratory, Bar Harbor, ME). Sox2-GT mice were generated by backcrossing Sox2/Cre (Hayashi S, 2002) and GCaMP5-tdTomato mice (Jackson Laboratory) resulting in ubiquitous expression of the same genetically encoded calcium reporters. MD-GFP mice on C57BL6/J background were generated by intercrossing nNOS/CreERT2 and mTmG/fl mice as described previously (both from Jackson Laboratory) (Riquier-Brison et al., 2018). Tamoxifen was administered 75 mg/kg by oral gavage for a total of three times (every other day) for full induction, and one time for partial induction (Gyarmati et al., 2021). NGF-eGFP reporter mice which express eGFP under the control of the mouse NGF promoter were generated and reported earlier (Kawaja et al., 2011). Some mice received low-salt diet (TD 90228, Harlan Teklad, Madison, WI) or chronic nerve growth factor (NGF 0.03 mg/kg in 100 µL PBS sc., N8133, Millipore Sigma, Burlington, MA) treatment for 10 to 14 days or, intravenous bolus arginine vasopressin (AVP, 5µg/kg, V9879 Sigma Aldrich, St. Louis, MO), isoproterenol (Iso, 10mg/kg I6504 Sigma Aldrich, St. Louis, MO), Angiotensin II (AngII, 800ng/kg, EK-002-12CE, Phoenix Pharmaceuticals, Burlingame, CA), Furosemide (Furo 1mg/kg, Sanofi-Aventis, Budapest, HU), gastrin (1µg/kg, Sigma Aldrich, St. Louis, MO), L-glutamic acid (0.75mg/kg, G1251, Sigma Aldrich, St. Louis, MO), Gamma-aminobutyric acid (GABA 50mg/kg, A2129, Sigma Aldrich, St Louis, MO), nerve growth factor (NGF, 1mg/kg, N8133, Millipore Sigma, Burlington MA), LM11a 31 dihydrochloride (LM11a, 25mg/kg, Tocris, Minneapolis, MN), or amyloid precursor protein (0.6 mg/kg, sAPPα, S9564 Sigma Aldrich Solutions, St. Louis, MO) injection. Some mice received low dose streptozocin treatment (50ug/g body weight once a day for five consecutive days, S0130, Sigma Aldrich, St Louis, MO) to induce diabetes mellitus. Animals with blood glucose levels over 300mg/dL were considered diabetic. All animal protocols were approved by the Institutional Animal Care and Use Committee at the University of Southern California.

### Intravital imaging using multiphoton microscopy (MPM)

Under continuous anesthesia (Isoflurane 1-2% inhalant via nosecone), mice, in which the left kidney was exteriorized through a flank incision, were placed on the stage of the inverted microscope with the exposed kidney mounted in a coverslip-bottomed chamber bathed in normal saline as described previously (Hackl et al., 2013; Kang et al., 2006). Body temperature was maintained with a homeothermic blanket system (Harvard Apparatus). Alexa Fluor 680-conjugated albumin (Thermo Fisher, Waltham, MA) was administered iv. by retro-orbital injections to label the circulating plasma (30 µL iv. bolus from 10 µg/ml stock solution).

The images were acquired using a Leica SP8 DIVE multiphoton confocal fluorescence imaging system with a 63× Leica glycerine-immersion objective (numerical aperture (NA) 1.3) powered by a Chameleon Discovery laser at 960 nm (Coherent, Santa Clara, CA) and a DMI8 inverted microscope’s external Leica 4Tune spectral hybrid detectors (emission at 510-530 nm for GCaMP5 and at 580-600 nm for tdTomato) (Leica Microsystems, Heidelberg, Germany). The potential toxicity of laser excitation and fluorescence to the cells was minimized by using low laser power and high scan speeds to keep total laser exposure as minimal as possible. The usual image acquisition (12-bit, 512×512 pixel) consisted of only one z stack per tissue volume (<1 min), which resulted in no apparent cell injury. Fluorescence images were collected in time series (xyt, 526 ms per frame) with the Leica LAS X imaging software and using the same instrument settings (laser power, offset, gain of both detector channels). The strong MD cell-specific tdTomato fluorescence signal (Shroff et al., 2021) and high-resolution MPM imaging allowed for easy identification of single MD cell bodies and processes.

### Quantification of GCaMP5 fluorescence intensity

An optical section including the vascular pole and the MD of a superficial glomerulus was selected, and time (xyt) series with 1 frame per 526 millisecond were recorded for 3 to 10 minutes to measure MD cell calcium dynamics. The strong, positive signal (GCaMP5/tdTomato fluorescence) and high-resolution MPM imaging allowed for easy identification of single MD, JG, EGM, IGM, EA and AA smooth muscle cell bodies. For the quantification of changes in mean GCaMP5 fluorescence intensity ROIs were drawn closely over the total cell body of single cells or over the entire MD plaque and the changes in GCaMP5 F/F_0_ (green channel; fluorescence intensity expressed as a ratio relative to baseline) were measured after the experiment in the defined ROI using the Quantify package of LAS X software (version 3.6.0.20104; Leica-Microsystems). MD whole plaque data were smoothed using exponential 2^nd^ order smoothing method (50 neighbors, GraphPad Prism). The frequency of MD calcium transients was measured based on the same defined ROIs using 3 to 10 minutes long time series recordings before and after manipulation. Maximum projection images of time series were generated (t=3min) to measure cumulative GCaMP5 fluorescence intensity over time.

### Pearson’s (*R*-based) connectivity analyses

Correlation analyses between the calcium signal time series for all MD cell pairs were performed with GraphPad Prism 9.0.1. as described (Salem et al., 2019). The correlation function R between all possible cell-pair combinations was assessed using Pearson’s correlation. Data were displayed as heat-map matrices, indicating individual cell-pair connections on each axis (minimum = −1; maximum = 1). PL<0.001 was deemed a statistically significant cell–cell connection. The coordinates of the imaged cells were used in the construction of connectivity line maps. Cell pairs (RL>0.35 and PL<0.001) were connected with a red straight line. MD cells with the most connections were labeled light red, MD cells without any apparent correlation to other MD cells were labeled light blue.

### Measurement of afferent renal nerve activity (ARNA)

Rats were anesthetized with isoflurane (2-3% in 100% O_2_) and prepared for renal afferent nerve recordings are described previously (DeLalio and Stocker, 2021a, b). Briefly, femoral artery and venous catheters (PE50 tubing) were implanted to monitor arterial blood pressure and infuse drugs, respectively. Animals were artificially ventilated via tracheotomy to maintain end-expiratory CO_2_ between 3.5 – 4.5% and O_2_ between 35 – 45% (Gemini Respiratory Gas Analyzer, CWE Inc). Body temperature was continuously monitored and maintained at 37±0.2°C using a rectal thermometer and circulating water pad. Through a retroperitoneal incision, the right adrenal artery was cannulated with heat stretched Micro-renthane® tubing. Then, the right renal nerve was placed on bipolar stainless-steel electrodes, insulated with KWIK-SIL, and sectioned proximally to the recording electrode to isolate ARNA. Nerves signals were filtered (300-1000Hz), amplified (10,000), and digitized (2000Hz) using a Micro1401 and Spike 2 software. The nerve signal was calculated by subtracting the noise obtained after sectioning the distal nerve-end. Values were normalized to baseline values set at 100%. Peak responses (10s) were compared to a 60-s baseline segment. After all surgical procedures were completed, isoflurane anesthesia was replaced by Inactin (120 mg/kg, 0.2ml/min, IV; Sigma T133). Experiments began at least 45 min later.

Intra-renal artery infusion of arginine vasopressin (AVP, 0.01-1.0 ug/mL, 50uL), isoproterenol (1000 ug/mL, 50uL) or saline was infused (15s) via the adrenal catheter. Solutions were flushed with 0.12mL of saline at 0.4mL/min. Infusions were separated by a minimum of 5 min. To control for the changes in arterial blood pressure as a stimulus for renal afferent nerve activity, arginine vasopressin (0.25ug/mL, 50uL) or sodium nitroprusside (100 ug/mL, 50uL) were infused intravenously.

### Tissue processing, histology and immunofluorescence

Immunofluorescence detection of proteins was performed as described previously (Riquier-Brison et al., 2018). Briefly, kidneys were perfused and fixed in 4% PFA for 2 hours at room temperature, embedded in paraffin, and sectioned 8 µm thick. Cells were cultured on sterile glass coverslips. After 14 days of differentiation cells were fixed in 4% PFA for 15 minutes at room temperature. For immunofluorescence analysis of antibody stains, slides were washed in 1XPBS. For antigen retrieval, heat-induced epitope retrieval with Sodium Citrate buffer (pH 6.0) or Tris-EDTA (pH 9.0) was applied. To reduce non-specific binding, sections were blocked with normal serum (1:20). Primary and secondary antibodies were applied sequentially overnight at 4° C and 2 hours at room temperature. Primary antibodies and dilutions were as follows: anti-tyrosine-hydroxylase antibody (TH 1:100, AB152, Millipore Sigma, Burlington, MA), anti-calcitonin gene related peptide antibody (CGRP:100, AB36001, Abcam Cambridge, UK), anti-synaptophysin antibody (SYP 1:100, PA5-27286, Thermo Fisher, Waltham, MA), anti-nerve growth factor receptor antibody (p75NTR 1:100, ab227509, Abcam Cambridge, UK), anti-tropomyosin kinase A/B antibody ( TRKA/B 1:100, A7H6R, Cell Signaling Technology Danvers, MA), anti-cyclooxygenase 2 (COX2) antibody (1:100, 12282S, Cell Signaling Technology, Danvers, MA,), anti-Na-K-2Cl cotransporter antibody (NKCC2 1:100, Developmental Studies Hybridoma Bank, created by the NICHD of the NIH and maintained at The University of Iowa, Department of Biology, Iowa City, IA 52242), anti-renin antibody (1:100, AF4277, R&D Systems, Minneapolis, MN), anti-phospho-Tau antibody (pTau^S199^1:100, 44-734G, Waltham, MA), anti-GFP antibody (GFP 1:200 Thermo Fisher Scientific, Waltham, MA). Alexa Fluor 488, 594, and 647-conjugated secondary antibodies were purchased from Invitrogen (Waltham, MA). Slides were mounted by using DAPI-containing mounting media (VectaShield, Vector Laboratories Inc., Burlingame, CA). Sections were examined with Leica TCS SP8 (Leica Microsystems, Wetzlar, Germany) confocal/multiphoton laser scanning microscope systems as described previously (Gyarmati et al., 2021; Riquier-Brison et al., 2018). For histological analysis Picrosirius red (PSR) staining was performed on mouse kidney sections using Sirius Red F3B (Sigma-Aldrich, St. Louis, MS). Images were visualized at 25× magnification using Leica TCS SP8 (Leica Microsystems, Wetzlar, Germany). Imaging software (Image J, National Institutes of Health and the Laboratory for Optical and Computational Instrumentation (LOCI, University of Wisconsin)) was used to calculate the percent area of red-stained collagen as described before (Ranjit et al., 2016).

### Tissue CLARITY

Three-dimensional imaging was performed as previously described (Lindström et al., 2018) by carrying out whole-mount immunofluorescence stains on slices of MD-GFP mouse kidneys. Slices were fixed in 4% formaldehyde in 1x phosphate buffer saline (PBS) at room temperature for 45 min, washed in 1XPBS, blocked in 1xPBS with 0.1% TritonX100 and 2% SEA Block (ThermoFisher Scientific) for 1 hour, and sequentially incubated in primary and secondary antibodies over 2 days. Primary antibodies were as follows: tyrosine-hydroxylase (AB152, MilliporeSigma, 1:100), GFP (ThermoFisher, 1:200). To clear tissue slices, the slices were dehydrated in methanol via increasing concentrations 50%, 75%, 100% diluted in PBS - each for 1hr - and subsequently submerged in a 50:50 benzyl benzoate/benzyl alcohol (BABB): methanol solution, followed by 100% BABB. High resolution imaging of MD plaques and the adjacent glomeruli was performed on a Leica SP8 multiphoton microscope using a 63X glycerol immersion objective.

### MD cell isolation

MD-GFP cells were isolated as described before (Gyarmati et al., 2021). Briefly, MD-GFP mice were anesthetized and perfused through the left ventricle with ice cold PBS, and the kidneys were harvested. Kidney cortex was isolated and digested using Hyaluronidase and Liberase TM enzyme combination (concentration: 2mg/mL and 2.5mg/mL respectively, from Sigma). After digestion, MD cells were isolated based on their genetic reporter expression (GFP) by using FACS ARIAII cell sorter and excitation wavelengths 488 and 633nm in sterile conditions. The highest tdTomato expressing cells were collected as controls representing distal tubule segments.

### RNA seq and bioinformatics

Whole-Transcriptome RNAseq was performed at the USC Norris Molecular Genomics Core. Cells were extracted using Qiagen miRNeasy purification kit following manufacturer’s protocol for total RNA purification (Qiagen cat#217004). Libraries were simultaneously prepared using Takara’s SMARTer Stranded Total-RNA Pico v2 library preparation kit following manufacturer’s protocol (Takara cat#634412). Prepared libraries were sequenced on Illumina Nextseq500 at 2×75cycles.

RNA-seq data was analyzed using the RNA-seq workflow in Partek Flow software (V10.1.21., Partek Inc., St. Louis, MO, USA). Briefly, the raw sequencing reads were first trimmed based on the quality score (Phred QC>=20, min read length=25 nt) before mapped to mouse genome build mm10 using Star 2.61 (Dobin et al., 2013) with default parameter settings and Gencode M21 mouse transcriptome annotation (Mudge and Harrow, 2015) as guidance. Gencode M21 was then used to quantify the aligned reads to genes using Partek E/M method. Finally, gene level read counts in all samples were normalized using Upper Quartile normalization (Bullard et al., 2010) and subjected to differential expression analysis using Partek Gene Specific Analysis method (genes with fewer than 10 aligned reads in any sample among a data set were excluded). The differentially expressed gene (DEG) lists were generated using the cutoff of FDR<0.05 and fold changes greater than 2.0 either direction. The top 50 DEGs based on FC was used to generate the tissue specific enrichment analysis using TissueEnrich (Jain and Tuteja, 2019) or GTEx. Gene set enrichment analysis of DEG (Partek Flow) was used to identify MD cell specific Gene Ontology (GO) terms that best describe biological processes in neurons. The normalized gene level read counts from the top 50 neuron-specific MD cell enriched genes in 6 GO term categories were used to generate the heatmap in Partek Genomic Suite (v7. Partek Inc., St. Louis MO USA).

Single cell RNA sequencing was prepared using 10x Genomics 3’ v3.1 (cat# 1000092) following manufacturer’s protocol. Samples were parsed into single cells using 10x Genomics Chromium Controller and libraries were simultaneously prepared. Prepared single cell RNA sequencing libraries were sequenced on the Illumina Novaseq6000 platform at a read length of 28×90 and read depth of 100,000 reads/cell for 2000-4000 cells. scRNAseq data was analyzed using the scRNAseq workflow by Partek Flow. Briefly, the raw sequencing reads with adaptors trimmed were mapped mm10 genome using Star 2.6.1 and quantified using Gencode M25 annotation to generate gene level counts. The gene counts were subjected to QA/QC and the low-quality cells were filtered using the following criteria: (1) contained less than 300 or more than 8000 detected genes, (2) mitochondrial counts higher than 15% of total counts. The counts were normalized using the Partek Flow recommended method (divided by 1 million, Add:1 and log2). Dimension reduction was carried out using PCA, followed by Graph-based Clustering with default settings and UMAP visualization (McInnes et al., 2018). Cell populations were determined by expression of relevant biomarkers. A 5-fold thresholding in *Nos1* and *Pappa2* expression was applied to filter out potential non-MD cell contamination. Nos1/Pappa2 expression filtered cells went through PCA, Graph-based clustering and UMAP visualization.

### Generation of the MD^geo^ cell line

Freshly isolated MD cells were plated at a density of 0.5^x^10^5^ cells/well in a 24-well plate and primary cultured at 37 °C 5% CO_2_ in Dulbecco’s Modified Eagle Medium: Nutrient Mixture F-12 (DMEM-F12, Gibco, Thermo Fisher Scientific) supplemented with 10% Fetal Bovine Serum (FBS, Thermo Fisher Scientific), 1% Penicillin-Streptomycin (P/S 10,000 U/mL, Thermo Fisher Scientific) and 0.0005% of Dexamethasone. At 80% confluence, cells were infected using Lenti-SV40 (ts58 temperature sensitive mutant, 10^6^ IU/mL, LV629, Applied Biological Materials) Lentivirus to achieve temperature sensitive immortalization, proliferation of MD cells at 33 °C and differentiation of MD cells at 37 °C according to manufacturer’s instructions. Briefly, cells were infected two times 8 hours apart using MOI:3 to achieve optimal viral density and incubated for 24 hours at 37°C, 5% CO_2_ in the presence of Polybrene (5ug/mL). Lentiviral vector was diluted with complete MD cell culture media to avoid cytotoxicity. After 24 hours incubation cells were cultured in complete MD cell culture media supplemented with INF-Gamma (0.01ug/mL) and nerve growth factor (NGF, 0.1 ug/mL; N8133, Millipore Sigma) at 33°C for proliferation. INF-Gamma concentration was decreased gradually after 1 week. Importantly, NGF supplementation was crucial for MD cell survival. Cells were subcultured after 48 hours. For differentiation, cells were incubated at 37°C, 5% CO2 for 14 days in complete MD cell culture media supplemented with NGF (0.1ug/mL; N8133, Millipore Sigma). All experiments were completed between 10th and 15th cell passage.

### Western Blot

Fully differentiated and undifferentiated MD^geo^ cells and MMDD1 cells (Yang et al., 2000) were incubated in normal salt (control), low salt, and NGF supplemented FBS-free culture media for 4 hours. Cells were starved of NGF for 48 hours and FBS for 12 hours before all experiments. Isosmotic normal and low salt media was made as described previously (Yang et al., 2000). Isosmotic normal salt media was supplemented with NGF as described above. Cells were harvested and used for Western blot analysis. Samples were separated on a 4–20% SDS-PAGE and transferred onto polyvinylidene difluoride membrane. The blots were blocked for 1 h with Odyssey Blocking Buffer (LI-COR Biosciences, Lincoln, NE) at room temperature. This was followed by an incubation of primary antibodies overnight at 4°C in 20 ml of Odyssey Blocking Buffer (LI-COR Biosciences), and 20 µl of Tween-20 (Sigma-Aldrich, St. Louis, MO) solution. Primary antibodies and dilutions were as follows: anti-p75NTR antibody (1:1000, ab227509, Abcam, Cambridge, UK), anti-COX2 antibody (1:1000 12282S, Cell Signaling Technology, Danvers, MA), anti-NKCC2 cotransporter antibody (1:1000 Developmental Studies Hybridoma Bank, created by the NICHD of the NIH and maintained at The University of Iowa, Department of Biology, Iowa City, IA 52242), anti-neuronal nitric oxide synthase antibody (Nos1 1:1000, SC648, Santa Cruz Biotechnology, Dallas, TX) anti-phospho and total extracellular signal-regulated kinase antibody (p/total ERK 1-2 1:1000 CST4396, CST4696, Cell Signaling Technology, Danvers, MA), anti-phospho and total protein kinase B antibody (AKT 1:1000, CST2920, CST4060, Cell Signaling Technology, Danvers, MA), anti-phospho and total IkB kinase antibody (p/total IkB 1:1000, 1:500, CST9246S, CST924, Cell Signaling Technology, Danvers, MA), anti-phospho and total panTrk (1:1000, 4619S, 92991S, Cell Signaling Technology, Danvers, MA), anti-renal outer medullary potassium channel antibody (ROMK 1:2000, APC-001, Alomone Labs, Jerusalem, Israel), anti-phospho and total p38 mitogen-activated protein kinase antibody (p/total p38 1:1000 Cell Signaling Technology, Danvers, MA). After incubation, blots were washed with PBS-T (1× PBS plus 1:1000 Tween-20) and then incubated in 20 ml of PBS, 20 ml of blocking buffer, 20 µl of Tween 20, and 20 µl of 10% SDS with goat anti-rabbit and goat anti-mouse secondary antibodies (1:15,000; LI-COR Biosciences). Blots were then visualized with Odyssey Infrared Imaging System (LI-COR Biosciences).

For immunoblotting of mouse cortical homogenates, manually dissected slices of kidney cortex were homogenized in a buffer containing 20 mM Tris·HCl, 1 mM EGTA pH 7.0, and a protease inhibitor cocktail (BD Bioscience, San Jose, CA). Protein (40 µg) was processed for immunoblotting as described above, using anti-p75NTR antibody (1:1000, ab227509, Abcam, Cambridge, UK) primary antibody.

### NO assay

To assess nitric oxide (NO) synthesis in MD^geo^ cells NO sensitive Diaminofluorescein-FM diacetate (D1946, DAF-FM DA, Millipore Sigma, Burlington, MA) was used as described before (Kovács et al., 2003). Briefly, fully differentiated MD^geo^ cells were loaded with DAF-FM DA (10µg/mL for 10 minutes) either in the presence or absence of the selective inhibitor of neuronal nitric oxide synthase (7-Nitroindazole (7NI) or Nω-propyl-l-arginine (NPA)). NO synthesis was measured based on the increase in the fluorescence intensity of DAF-FM DA (F/F_0_ green channel; fluorescence intensity expressed as a ratio relative to baseline) over time. Fluorescence emission was detected at baseline, and after 30, 60, and 90 minutes at 515 +/− 15 nm emission wavelength in response to 495 nm excitation using Leica SP8 DIVE multiphoton confocal fluorescence imaging system. Changes in DAF-FM DA F/F_0_ were measured after the experiment using the Quantify package of LAS X software (3.6.0.20104; Leica-Microsystems)

### PGE2 biosensor technique

PGE2 biosensor cells were used as described before (Peti-Peterdi et al., 2003). In brief, PGE2 biosensor cells were specifically engineered human embryonic kidney cells (HEK 293 cells) to express the calcium-coupled PGE2 receptor EP1, leading to a calcium response upon PGE2 binding. HEK-EP1 cells were loaded with Fluo-4 and Fura Red (1 μM, for 10 min) and positioned next to the fully differentiated MD^geo^ cells in culture. MD^geo^ cell PGE2 production was measured based on the biosensor cell intracellular Ca^2+^ concentration signal, which was detected by increases in Fluo-4/Fura Red (F/F_0_) fluorescence intensity ratio over time. Fluorescence emission was detected every 526 ms using a Leica SP8 DIVE multiphoton confocal fluorescence imaging system as described above (emission at 460-520 nm for Fluo-4 and at 580-640 nm for Fura Red) (Leica Microsystems, Heidelberg, Germany).

### Glomerular Filtration Rate measurement

GFR measurements were performed using the MediBeacon Transdermal Mini GFR Measurement System (MediBeacon) as described previously (Scarfe et al., 2018). Briefly, mice were anesthetized and the MediBeacon sensor was placed on the depilated dorsal skin. Mice were injected retro-orbitally with the inulin analog exogenous GFR tracer fluorescein-isothiocyanate conjugated sinistrin (FITC-S 7.5 mg/100 g body weight, MediBeacon, St. Louis, MO). The excretion kinetics of the FITC-S was measured for 90 minutes. GFR was then calculated based on the decay kinetics (half-life time) of FITC-S using MediBeacon Data Studio software (MediBeacon).

### Statistical methods

Data are expressed as average ± SEM and were analyzed using Student’s t-tests (between two groups), or ANOVA (for multiple groups) with post-hoc comparison by Bonferroni test. P<0.05 was considered significant. Statistical analyses were performed using GraphPad Prism 9.0.1 (GraphPad Software, Inc.).

## ACKNOWLEDGMENTS

This work was supported in part by US National Institutes of Health grants DK064324, DK123564, and S10OD021833 to J.P-P and R01 HL152680 to S.D.S. U.N.S. was funded by predoctoral research fellowship 19PRE34380886 of the American Heart Association. The Genotype-Tissue Expression (GTEx) Project was supported by the Common Fund of the Office of the Director of the National Institutes of Health, and by NCI, NHGRI, NHLBI, NIDA, NIMH, and NINDS. The data used for the analyses described in this manuscript were obtained from the GTEx Portal on 07/15/2020.

## AUTHOR CONTRIBUTIONS

G.G. and J.P.P. designed the study, analyzed the imaging data, and wrote the manuscript. U.N.S, A.R.B, A.I., S. D., D. B., A.W. J., L. M. and B. V. Z. made substantial contributions to acquire data. Y.C. filtered and analyzed transcriptome data. All authors approved the final version of the manuscript.

## DECLARATION OF INTERESTS

J.P-P. and G.G. are co-founders of Macula Densa Cell LLC, a biotechnology company that develops therapeutics to target macula densa cells for a regenerative treatment for chronic kidney disease. Macula Densa Cell LLC has a patent entitled “Targeting macula densa cells as a new therapeutic approach for kidney disease” (US patent 10,828,374).

**Supplemental Figure S1. Additional features of Sox2-GT and MD-GT mice.**

**(A)** Illustration of the experimental workflow using transgenic mouse models, Cre-lox induction strategies, and intravital MPM imaging of cell calcium over time to study 4D physiology in comparative (cc4DP), multi-cell (mc4DP), and single-cell (sc4DP) modes. Representative intravital MPM images are shown for each of the three experimental modes.

**(B)** Intravital imaging of macula densa (MD) cell calcium in cc4DP mode. Left: Representative recordings and MPM image of GCaMP5 fluorescence intensity normalized to baseline (F/F_0_) in proximal tubule (PT, blue), MD (green), and distal tubule (DT, magenta) cells in Sox2-GT mice. Center: Distribution of single MD cells (shown as fraction of total number of cells) based on their cumulative GCaMP5 fluorescence intensity in maximum projection images (MPI) of 2-min time lapse recordings and based on their Ca^2+^ firing frequency.

**(C)** Left: Representative recordings of normalized GCaMP5 fluorescence intensity of a single MD cell in vivo in the intact mouse kidney or in vitro after isolation from freshly digested kidneys of MD-GT mice. Right: Overlay of the Ca^2+^ recordings of all 21 individual MD cells shown in Fig. 1B. Note the clustering of single-cell Ca^2+^ transients and their periodic oscillations over time (curve fitting in green).

**Supplemental Figure S2. Initial characterization and validation of the MD cell transcriptome.**

**(A)** Main features of the MD cell transcriptome. Left: 3D principal component analysis (PCA) showing clustering of the MD cell versus control cell transcriptome (n=4 MD and n=8 control cell samples). Right: Volcano-plot of up-regulated (red) and down-regulated genes (green) in MD vs control cells based on values of false discovery rate (FDR). Note the higher number of down-regulated compared to up-regulated genes.

**(B)** Validation of the specificity of MD cell-derived transcriptome showing the high-level expression of classic MD-specific genes in all four MD samples, including Nos1, Slc12a1, Nt5c1a, and Avpr1a.

**(C)** Tissue specificity analysis of the control kidney cell transcriptome. Bar chart showing the enrichment (−Log10(P-Adjusted)) of the top 50 highest expressed control cell-specific genes in various tissues using TissueEnrich and the Mouse ENCODE dataset for comparison.

**Supplemental Figure S3. Validation and characterization of the new MD cell line MD^geo^.**

**(A)** Representative immunofluorescence images of Cox2 and Nkcc2 labeling in mMD^geo^ cells showing cytosolic or membrane-targeted localization, respectively. Negative control shows endogenous genetic mGFP expression only. Nuclei are labeled blue with DAPI. Note the co-localization of mGFP and Nkcc2 signals (yellow in overlay) in the cell membrane. In addition to the overlay, separate red and green channels are shown at the bottom.

**(B)** Representative immunoblots and statistical summaries of Cox2, Nkcc2, and Romk expression in mMD^geo^ cells cultured at 37° and 33°C, and in the previously established but now extinct MMDD1 cell line using immunoblotting of whole cell lysates. The effect of low salt (LS) versus normal salt (NS) culture condition on the ratio of phospho/total ERK1/2, COX2, and phospho/total p38 in differentiated (37° C) mMD^geo^ cells. Ns: not significant, *: p<0.05, ****: p<0.0001, data are mean ± SEM, n=4-5 each.

**(C)** Measurement of nitric oxide (NO, top row) and PGE_2_ (bottom row) synthesis and release in MD^geo^ cells in timed control (at baseline and 60 min) and after preincubation with either selective Nos1 inhibitor NO-propyl-L-arginine (NPA, 300 mM) or selective COX-2 inhibitor SC58236 (100 nM) for Nos1-mediated NO and Cox2-mediated PGE2 detection, respectively. For NO measurement, cells were loaded with the fluorescent NO indicator DAF-FM at 37 °C, and changes in DAF-FM fluorescence intensity (green) were monitored using time-lapse confocal microscopy. Maximum change in DAF-FM fluorescence intensity in the cytoplasm of MD cells was evaluated after 60 minutes and compared to baseline. Note the high DAF-FM fluorescence intensity (heterogenous pattern) in control conditions indicating NO synthesis in MD^geo^ cells. Summary of normalized DAF-FM fluorescence intensity in control condition and after NPA treatment (right panel, n=4 wells, 25 measurements each). For the detection of MD^geo^ cell PGE_2_ release, representative maximum projection images of the HEK/EP1 PGE_2_ biosensor cell signal (cell Ca^2+^ detected by Fluo4) are shown at baseline and over time. Note the high HEK/EP1 PGE_2_ biosensor cell signal in control conditions indicating PGE_2_ release from MD^geo^ cells. Summary of HEK/EP1 PGE_2_ biosensor cell signal (normalized Fluo4 fluorescence) in control and after selective COX-2 inhibition ( COX2i). ***: p<0.001, ****: p<0.0001, data are mean ± SEM, data points represent the average of ten measurements in n=10 wells each.

