## Supplementary figures and images for "Neuron-like function of the nephron central command"

### Supplemental Figures 1-3

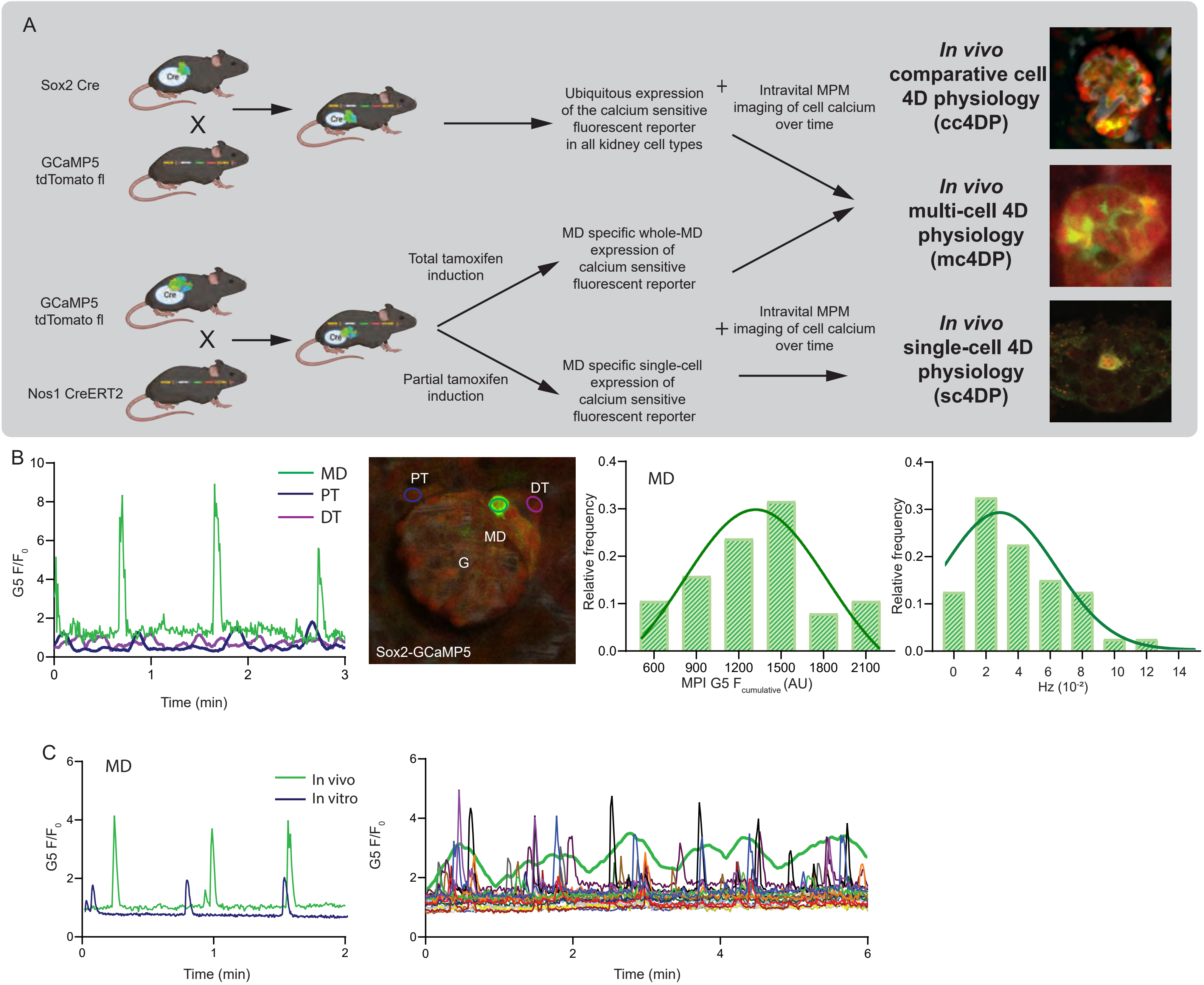

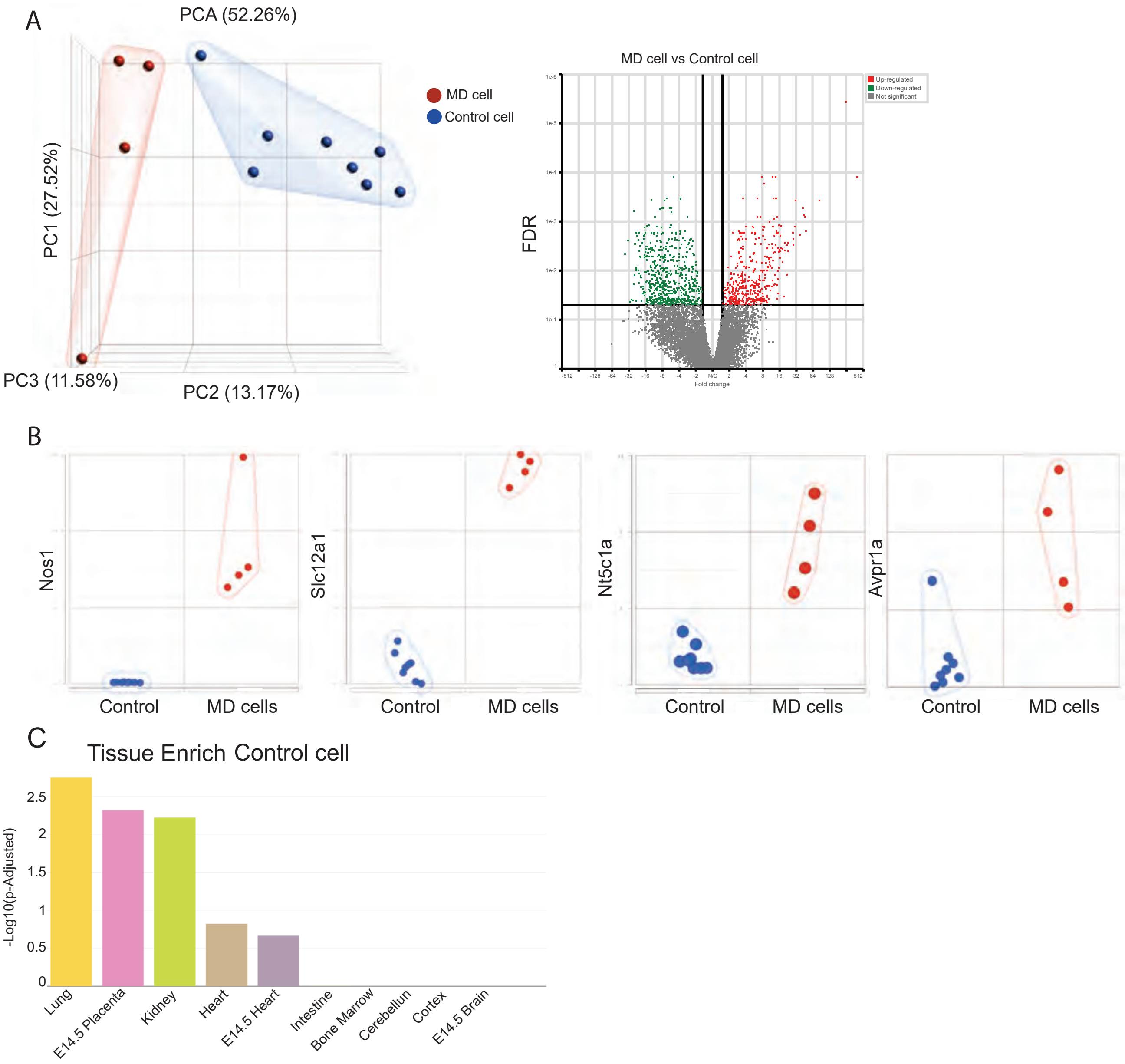

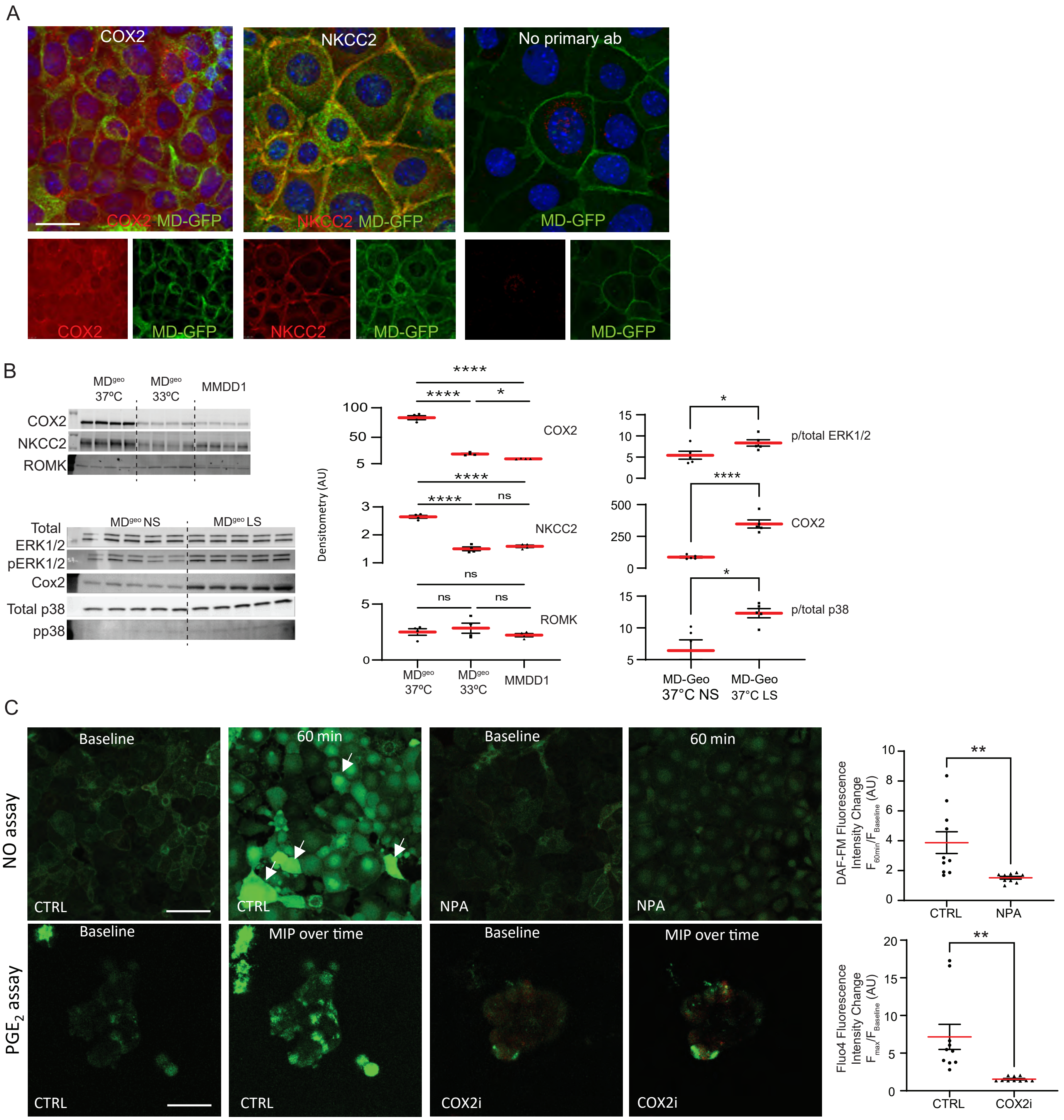
